# distQTL: Distribution Quantitative Trait Loci Identification by Population-Scale Single-Cell Data

**DOI:** 10.1101/2025.04.04.647121

**Authors:** Alexander Coulter, Chun Yip Tong, Yang Ni, Yuchao Jiang

## Abstract

Mapping expression quantitative trait loci (eQTLs) is a powerful method to study how genetic variation influences gene expression. Traditional bulk eQTL methods rely on averaged gene expression across a possibly heterogeneous mixture of cells, which can obscure underlying regulatory heterogeneity. Single-cell eQTL methods circumvent the averaging artifacts, providing an immense opportunity to interrogate transcriptional regulation at a much finer resolution. Recent developments in metric space regression methods allow the use of full empirical distributions as response objects instead of simple summary statistics such as mean. Here, we leverage Fréchet regression to identify distribution QTLs (distQTLs) using population-scale single-cell RNA sequencing data. We apply distQTL to the OneK1K cohort, consisting of scRNA-seq data of peripheral blood mononuclear cells from 982 donors, and compare results to various eQTL approaches based on summary statistics and mixed effects modeling. We demonstrate the superior performance of distQTL across different gene expression contexts compared to other methods and benchmark our results against findings from the Genotype-Tissue Expression Project. Finally, we orthogonally validate calls from distQTL using cell-type-specific epigenomic profiles.

## Introduction

Expression quantitative trait loci (eQTLs) link single-nucleotide polymorphisms (SNPs) to gene expressions, offering insights into the molecular mechanisms of genetic variants that are associated with complex traits (1). High-throughput RNA sequencing (RNA-seq) data, paired with DNA genotype information, has achieved huge successes at eQTL-mapping, such as the identification of tissue-specific eQTLs from the Genotype-Tissue Expression (GTEx) Project (2). These studies, however, were at the bulk-tissue level, where one could only observe averaged expression across a possibly heterogeneous mixture of cells, and the true underlying cellular heterogeneity was attenuated.

Recent advancements in single-cell RNA-seq (scRNA-seq), along with its increasing accessibility and decreasing cost, have facilitated population-scale transcriptomic profiling at the cellular level, enabling the discovery of cell-type-specific eQTLs (3; 4; 5; 6). For instance, the OneK1K cohort (7) consists of scRNA-seq data of ∼1.27 million peripheral blood mononuclear cells (PBMCs) collected from 981 donors, resulting in the discovery of novel cell-type-specific eQTLs. See recent reviews (8; 9) and database (10) for existing scRNA-seq data utilized for eQTL discovery.

Existing eQTL studies for scRNA-seq, while revelatory, most commonly identify cell clusters and annotate cell types from the single-cell data first, and then calculate average expression per individual for each cell type by constructing pseudo-bulk samples (4; 11). This partially forfeits the purpose of single-cell sequencing, which goes beyond the mean measurements to characterize the distribution of gene expression. Although methods for analyzing higher-order distribution features of gene expression have been developed, including variance QTL (vQTL) detection (12; 13), they have been largely confined to bulk eQTL-mapping. One notable exception is (14), where zero-inflated negative binomial distributions were fit to donor-level scRNA-seq data, giving mean, variance, and over-dispersion estimates. However, in applied data analysis they found no vQTLs independent of mean QTLs, owing to the well-established mean-variance dependence in scRNA-seq data (15).

Beyond pseudo-bulk approaches, attempts to account for cell-cell correlations have been made with mixed-effects Poisson regression models (6). However, its high computational cost and non-convergence issue make it practically infeasible to scan every gene-SNP pair for data sets as large as the OneK1K cohort. Additionally, introducing an individual-specific random effect has been shown to result in a lack of power (4). Finally, the class of mixed-effects models are only concerned with mean effects, missing out on higher-order distribution features.

Non-mean effects have been suggested to indicate gene-environment interactions (16; 17) or gene-gene interactions (18), multi-allelic effects (19), and bursting kinetics (20; 21). Feedback models of gene expression are generally not mean-reducible (22; 23; 24), suggesting mean-only testing is insufficient for capturing all biologically relevant expression and other single-cell level phenotypic differences (24). Full gene expression distributional approaches have been applied to single-cell data, including single-cell probabilistic trait loci (scPTL) (24) and IDEAS model (25), where the latter can be viewed as a covariate-dependent extension of the former. However, both methods require calculating a donor-donor distance matrix per gene, which scales poorly to population-scale data sets such as the OneK1K cohort. Importantly, neither provides covariate-dependent distributional estimates, a significant omission in characterizing distributional effects.

In light of various sources of biological variability that can be neglected by mean-only tests, we propose a statistical method based on Fréchet regression (26) and associated asymptotic partial *F* -test (27; 28) to identify distribution QTLs (distQTLs). We directly use the empirical distribution of library-size corrected gene expression at the donor level as the response, performing both hypothesis testing and calculating conditional distributions under one coherent theory. Characterizing gene expression distribution is an omnibus approach that broadens inference beyond the first and second moments (i.e., mean and variance) from scRNA-seq, while capturing expression level, variability, and stochasticity in a unified manner (21; 20). distQTL-mapping also scales linearly with donor count and uses a fraction of the time required for Poisson mixed-effects regression (less than one-tenth of a second per model, contrasted to minutes), enabling its use in population-scale applications.

We compare distQTLs with baseline eQTL-mapping using mean (*µ*QTL), variance (vQTL), coefficient of variation (cvQTL), and bursting fraction (percentage of cells with nonzero expression, bQTL) as outcomes. We apply a correction to distQTL and baseline eQTL p-values to account for misspecified null distribution. We demonstrate our proposed framework in the OneK1K cohort, identifying more cell-type-specific eQTLs than the pseudo-bulk methods. We benchmark against existing reports to assess performance and further evaluate type I error control via permutation. Altogether, this work provides an atlas of how genetic variation influences gene expression across cell types with various statistical testing schemes that corroborate and complement existing studies and approaches. distQTL is an open-source R package available at https://github.com/alexandercoulter/distQTL.

## Materials and Methods

### QTL Identification

For each gene-SNP pair, we regress cell-type-specific gene expression against SNP genotype while controlling for a set of fixed donor-level covariates. We index donors by 1 ≤ *i* ≤ *n*, cell-type groups by 1 ≤ *k* ≤ *K*, individual donor and group level cells by 1 ≤ *c*_*i,k*_ ≤ *C*_*i,k*_, genes by 1 ≤ *g* ≤ *G*, and *cis*-SNPs by 1 ≤ *s*_*g*_ ≤ *S*_*g*_. Denote by 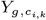 the raw gene expression for cell *c*_*i,k*_, i.e., from donor *i* and cell-type group *k*. Similarly, denote by 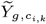 the library size corrected gene expression for cell *c*_*i,k*_. Let **x**_*i*_ be the fixed donor-level covariates. In our later OneK1K cohort data analysis, the covariates are quadratic orthogonal polynomials of age (`age` and age^2^), sex (`male` and `female`), and the top leading genotype principal components. Generally, the choice of covariates is context-dependent and up to the practitioner; see for instance the Discussion for details on additional covariates such as the probabilistic estimation of expression residuals (PEER) factors or gene expression PCA. Finally, let 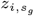 be the individual SNP genotype encoded as 0/1/2, also included as a covariate in all regression models. All regression methods are applied within cell-type group; in the following equations, the *k* subscript is dropped for simplicity.

#### Baseline and univariate eQTL

We calculate donor-level univariate summaries *f* ( · ) for library size corrected gene expression, fitting linear models

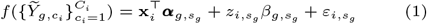

Where 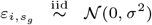. For each donor and gene, the univariate expression summaries include the sample mean, sample variance, proportion of cells with positive gene expression (“bursting”), and sample coefficient of variation (CV). For each expression summary, we test null hypothesis 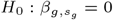 with a Wald *t*-test.

We perform p-value correction by a permutation method, permuting responses for each model once and obtaining a permutation p-value for each gene-SNP pair. We pool permutation p-values for each cell-type group to obtain empirical null-corrected p-values *p*_*c*_. This is similar to (29) and related R function `qvalue::empPvals` (R package version 2.38.0), though we leave very small p-values unadjusted; see Supplement (Section S2). Unless otherwise stated, we adopt a significance level of 10^−4^; practitioners may apply other *post hoc* correction methods for multiple testing. Hereafter, we refer to these four baseline eQTL-mapping methods as *µ*QTL, vQTL, bQTL, and cvQTL, respectively.

#### Poisson mixed-effects regression (PMER)

We model cell-level raw gene expression with PMER (6), fitting a random intercept per donor, utilizing the `glmer` function from the `lme4` R package (30). Library correction factors are used as regression offsets to account for differences in gene expression measurement exposure. We use the same fixed effects structure as eq. (1) and test null hypothesis 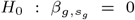 with a Wald *t*-test. Because the computation time of PMER is very high, we implement this procedure only on a subset of gene-SNP pairs, obtained by selecting the top eQTL identified by distQTL within B cells. We do not perform p-value correction by permutation for PMER results since our fixed effects design does not differ within-donor, and there are no meaningful permutations which preserve donor-level cell-cell correlation structure.

#### Distribution QTL (distQTL)

We implement distributional Fréchet regression (26) with gene- and donor-specific empirical distributions of 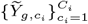. Unlike any of the aforementioned univariate summaries, Fréchet regression aims to model the entire distribution of cell-type-specific gene expression as a function of covariates and *cis*-SNP genotype. Fréchet regression generalizes the Euclidean regression problem, which takes covariate vector *X* ∈ ℝ^*p*^ (where 𝔼 (*X*) = 0 and Var(*X*) = Σ) and response *Y* ∈ ℝ and finds conditional mean

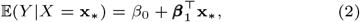

where *β*_0_ and ***β***_1_ are intercept and slope parameters. The least-squares solutions for *β*_0_ and ***β***_1_ are obtained by solving the normal equations,

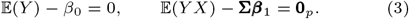

The space of distribution functions is not a vector space. As such, the linear model posed in eq. (2) cannot serve as the basis itself for regression with distribution responses. The observation of (26) is that eq. (2) can be reformulated without explicit dependence on these parameters, using solutions to eq. (3),

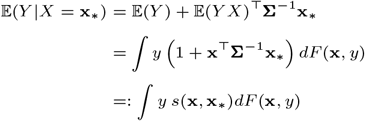

The final equation is a weighted expectation, where the weights *s*(**x, x**_∗_) := 1 + **x**^⊤^**Σ**^−1^**x**_∗_ satisfy ∫*s*(**x, x**_∗_)*dF* (**x**, *y*) = 1. The conditional mean therefore solves

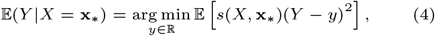

which is a weighted variance-minimizing problem. Eq. (4) is a function of only covariates, responses, and Euclidean distance. In the sample setting 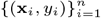 with plug-in estimator 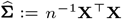, the sample predictors ŷ—the vector of estimated conditional means 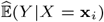—are obtained by projecting **y** onto the column space of (1, **X**).

Importantly, eq. (4) can be generalized to accommodate non-Euclidean responses, such as univariate distributions. We represent distributions in the space of univariate quantile functions 𝒬 := {**q** : (0, 1) ↦ ℝ, *u* ≤ *v* ⇒ **q**(*u*) ≤ **q**(*v*)}, and choose the 2-Wasserstein distance 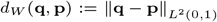. Then by substituting the Euclidean distance with the 2-Wasserstein distance in eq. (4), the Fréchet regression problem is

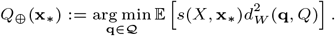

In the sample setting 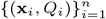 with plug-in estimators, the **x**_*i*_-conditional response estimate is the solution to

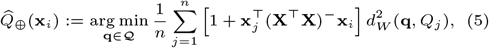

where **A**^−^ is any generalized inverse of **A**. In this paper, we refer to quantities from eq. (5) as *conditional mean distributions* or *conditional mean curves*; see (26) for more background. We comparatively illustrate hypothesis testing between linear and distributional regressions in Supplement (Section S1.4; Fig. S2).

To conduct distributional regression for population-scale scRNA-seq data, we calculate donor-specific empirical quantile functions (EQFs) per gene and cell-type group, using R function `quantile` (`type = 1` option), on a common *m*-grid in (0, 1). We use the same regressor set as in eq. (1), and test null hypothesis *H*_0_ : *Q*_⊕_(**x**, *z*) = *Q*_⊕_(**x**) with asymptotic partial *F*-test (27; 28); see Supplement (Section S1; Fig. S1). As with the univariate methods, we permute responses and refit once for each gene-SNP pair to perform cell-type-specific p-value correction as described in Supplement (Section S2). Hereafter, we refer to this eQTL-mapping method as *distQTL*.

### Simulations

We show the capacity of Fréchet regression to identify varying mean and non-mean effects through different illustrative examples utilizing zero-inflated negative binomial (ZINB) simulations. We generate covariate vectors 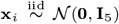 for *i* = 1, …, 100 samples/donors. For each sample, we draw *N* ∼ Poisson(200) values from a covariate-dependent ZINB distribution, analogous to cells. For simplicity, the parameters of the true underlying ZINB distribution are functions only of the first covariate *X*_1_, in three different effect type settings.

In setting A, we generate each true ZINB distribution such that the zero-inflation is fixed at zero (no inflation), the mean is a log-linear function of *X*_1_ with Gaussian errors, and the variance has log-normal sampling distribution independent of *X*_1_. In setting B, the zero-inflation is fixed at zero, the variance is a log-linear function of *X*_1_ with Gaussian errors, and the mean has log-normal sampling distribution independent of *X*_1_. In setting C, the mean and variance are sampled *iid* log-normal, the total positive proportion P[*Y >* 0] is a logistic function of *X*_1_ with Gaussian errors, and the size and probability parameters (*r, p*) of the NB component are chosen to satisfy the system of equations relating E(*Y* ), Var(*Y* ), and P[*Y >* 0]. We use six incrementally increasing *X*_1_ effect magnitudes, starting from zero, and perform 5,000 replicates for each combination of setting and effect size.

We calculate univariate summaries (sample mean, variance, % positive, and CV) for each sample-level group of “cells”, and perform Wald *t*-tests for *H*_0_ : *β*_1_ = 0. We evaluate sample-level EQFs on a common uniform *m* = 200 grid in (0, 1), and perform asymptotic partial *F* -tests for *H*_0_ : *Q*_⊕_(*X*) = *Q*_⊕_(*X*_−1_). To investigate the effect of under-discretization, we also calculate EQFs using *m* = 100 and *m* = 150; see Supplement (Fig. S3). For each *t*-test or *F* -test we record the corresponding − log_10_(*p*)-value.

### OneK1K Data and Quality Control

We use the OneK1K cohort (7) data set, consisting of scRNA-seq data of ∼1.27 million PBMCs collected from 981 donors of European ancestry, along with corresponding donor-level genotype data. Integer-valued scRNA-seq read count data across 36,571 genes and 1,248,980 cells are downloaded from the Human Cell Atlas (HCA) and corrected for library-size factor. Processed genotype data are provided by the authors of the OneK1K paper. For quality control, we remove: (i) doublets and cells with cell-type prediction scores *<* 0.6 or extreme total coverages (number RNA reads *>* 3 M.A.D. from median); (ii) donors with extreme cell numbers (*>* 3 M.A.D. from median); and (iii) SNPs with low rare allele presence (fewer than five donors with the minor allele). We further filter out each gene with no SNPs within 200Kbp from the gene’s transcription start site.

Cell-type labels are obtained from the HCA, and we focus on four major cell types: 102,179 B cells (naïve, memory, and transitional), 42,634 monocytes (CD14-positive, CD14-low, and CD16-positive), 221,802 CD4-positive T cells (central memory and effector memory), and 144,030 CD8-positive T cells (central memory and effector memory). Within each cell-type group, we filter out donors with fewer than 10 cells, and filter out genes with donor-level cell-type-specific nonzero expression rates below 1%. For fitting cell-type-specific EQFs, we choose *m* = 200, which exceeds or approximately matches median per-donor cell counts in these cell-type groups (B cells, 93; monocytes, 25; CD4+ T cells, 218; CD8+ T cells, 122).

### Benchmarking and Validation

eQTL findings are validated against the whole blood bulk eQTL results from the GTEx Project (2). We primarily benchmark OneK1K results using an aggregated cell group, to make the single-cell data comparable to the bulk data from GTEx. To reflect higher median per-donor cell count in aggregation (i.e. 520 cells per donor), we increase EQF discretization to *m* = 500. Since GTEx whole blood samples include granulocytes and other cell types not present in the PBMC-only samples in OneK1K, we also compare eQTL-mapping results in cell-type-specific groups to see how GTEx overlap is affected by relative cell-type abundance; see Supplement (Table S3). We illustrate eQTL-mapping overlap by contour plots, of proportion of OneK1K eQTL matches in GTEx against proportion of *cis*-SNPs that GTEx labels as eQTLs.

To further assess the performance of distQTL and to benchmark against the other univariate eQTL methods, we adopted cell-type-specific chromatin immunoprecipitation sequencing (ChIP-seq) data of histone modification in human blood cells (31). Specifically, we compared the ChIP-seq signals for activation and enhancer markers (H3K27ac, H3K4me1, H3K4me3) and repression marker (H3K27me3) at eQTLs identified by distQTL, at eQTLs uniquely detected by distQTL but not by the other methods, and at randomly selected SNPs serving as negative controls.

### Model Fit Run Time

We calculate total time to gene-SNP models in each cell-type group for distQTL and the competing eQTL-mapping methods. For all eQTL-mapping methods except PMER, we fit models to expression data from highly expressed genes in original ordering and with permuted responses, for approximately 5.8 million models across cell-type groups. For PMER, due to long per-model fit times, we select randomly 1000 genes with high expression in each cell-type group and fit one model per gene using the top eQTL identified by distQTL in B cells. All model fits were performed splitting genes in parallel over nine cores, and we estimate fit time per model using aggregate run time divided by number of models fit, multiplied by 9.

## Results

### Simulations

We simulate ZINB distributions in different covariate-dependent settings to demonstrate efficacy of distribution-based Fréchet regression. For simplicity, only *X*_1_ influences the responses.

In the first setting (Fig. 1**A**), only the mean is a function of *X*_1_. As expected, vQTL does not have statistical power in this setting. On the other hand, for non-vQTL methods we observe the distribution of − log_10_(*p*)-values shifts more positive as effect size increases, indicating increasing power with effect magnitude. In the second setting (Fig. 1**B**), only the variance is a function of *X*_1_. *µ*QTL does not have statistical power in this setting, while we observe non-*µ*QTL methods, including distQTL, exhibit increasing power as effect size increases. Finally, in the third setting (Fig. 1**C**), only the proportion of positive values is a function of *X*_1_. Neither *µ*QTL, vQTL, nor cvQTL exhibit statistical power in this setting, as expected; however, distQTL exhibits higher statistical power than bQTL. Across all settings, distribution based regression has comparable or superior power to univariate response regression methods, even in cases where certain univariate methods exhibit no sensitivity.

**Fig. 1.**
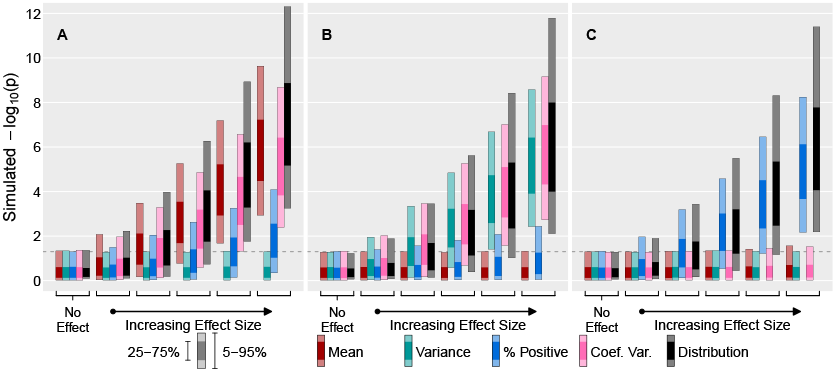
Comparisons of partial *F* -tests in simulated covariate-dependent, ZINB distributions, using different response targets. Each sample *i* = 1, …, 100 (i.e. donor) consists of *N* ∼ Poisson(200) ZINB draws (i.e. cells) on which univariate summaries and distributions are calculated. EQF discretization fixed at *m* = 200; covariates fixed at *p* = 5, where only *X*_1_ has varying effect size on (**A**) mean but not variance, (**B**) variance but not mean, and (**C**) positive proportion but neither mean nor variance. Results from 5000 iterations per setting.

### OneK1K eQTL Identification

#### distQTL v.s. pseudo-bulk and univariate eQTL

Fig. 2**A** illustrates donor-level univariate targets for POLE4 gene across levels of rs1861457 SNP genotype (ref/alt alleles G/A) in B cells, with associated *t*-test p-values. Fig. 2**B** shows donor-level EQFs and conditional distributions for the same gene-SNP pair, and Fig. 2**C** shows overlay of conditional distributions with Wasserstein partial *F* -test (i.e. distQTL) corrected p-value. Mean expression is not significantly associated with SNP genotype, but non-mean univariate responses are significantly associated with genotype, such as coefficient of variation (*p* = 1.13E−4). In Fréchet regression with distributional responses, the object of interest is the conditional distribution. distQTL, which evaluates the whole conditional distribution of POLE4 expression, identifies the most significant association (*p* = 1.97E−5) with SNP genotype. EQF results suggest distributions differ by higher-order moments (Fig. 2**C**); indeed, a further *t*-test with donor-level expression skewness as response has uncorrected p-value *p* = 1.72E−5.

**Fig. 2.**
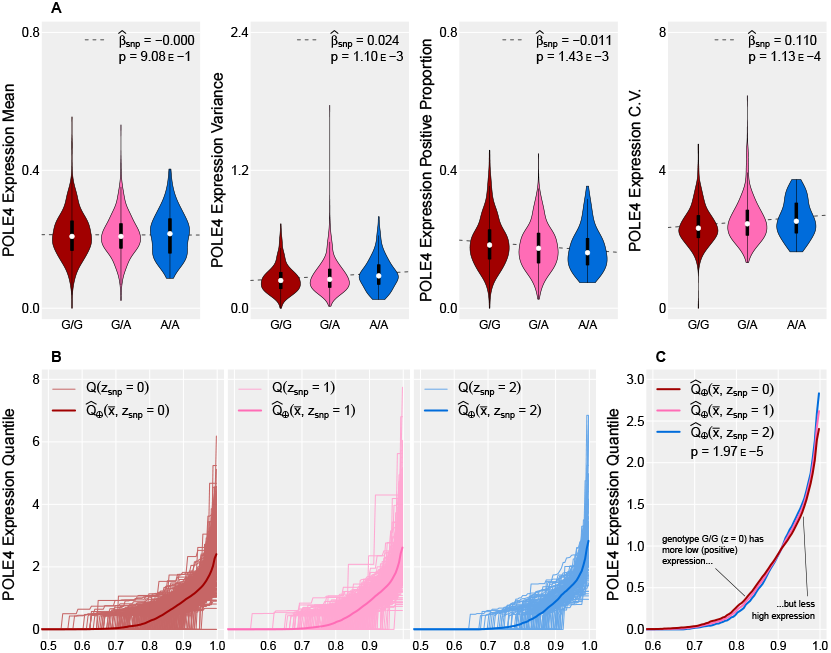
(**A**) Violin plots of donor-level B cell POLE4 gene expression summaries and partial *F* -test corrected p-values over levels of rs1861457 SNP. (**B**) Donor-level B cell POLE4 gene expression distributions (as EQFs) and conditional distributions, across levels (*z* = 0, 1, 2) of rs1861457 SNP. (**C**) Conditional distributions and Wasserstein partial *F* - test (i.e. distQTL) null-corrected p-value; note the different *x*-axis and *y*-axis scales.

Fig. 3**A** shows observed distQTL p-value distribution before and after p-value correction. The estimated null p-value distribution indicates uncorrected distQTL p-values exhibit inflated Type I error, which can be ameliorated by pooled permutation-based correction (29); see Supplement (Section S2). The distribution of corrected p-values is nearly flat over the support while retaining a slightly reduced spike toward lower values. This demonstrates that our correction method properly adjusts for the mis-specified null distributions. We observe similar correction patterns in other cell-type groups (Fig. S5). Quantile-quantile (QQ) plots (Fig. 3**B**) illustrate that, post p-value correction, distQTL is more powerful in detecting eQTLs in B cells than univariate summary regression methods, most comparable with *µ*QTL. The results can be reproduced in other cell-type groups (Fig. S5).

**Fig. 3.**
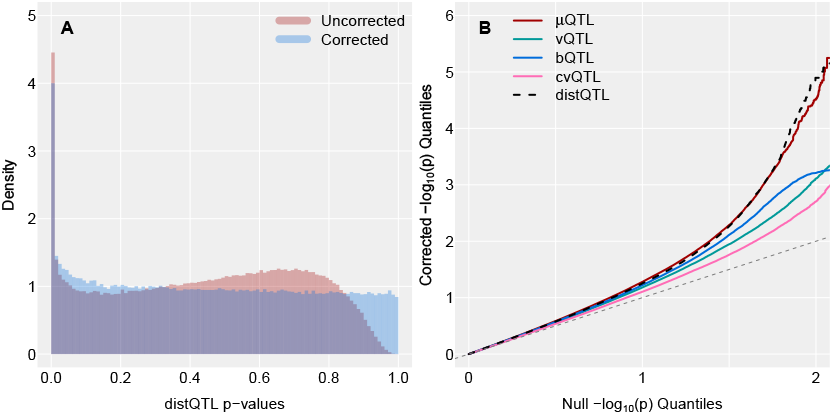
P-value correction and power calculation. (**A**) Empirical histograms of raw and null-corrected distQTL p-values from gene-SNP pair models in B cells. (**B**) QQ plot of null-corrected − log_10_(*p*)-values for gene-SNP pairs in B cells.

On the genome-wide scale, distQTL identifies more eQTLs when compared to the other methods, exemplified by the upset plots for the different cell types (Fig. 4; Fig. S6). In particular, distQTL identifies more gene-SNP pairs as strongly significant (*p* ≤ 10^−4^) in B cells while other methods label them as non-significant (*p >* 0.1), than vice versa – 4*/*0 compared to *µ*QTL, 163*/*0 compared to vQTL, 532*/*37 compared to bQTL, and 2409*/*25 compared to cvQTL. This pattern holds across all cell-type groups (Table S2). distQTL can be used to identify gene-SNP pairs that exhibit different expression patterns across cell types; a selection of such pairs are illustrated in Supplement (Fig. S4).

**Fig. 4.**
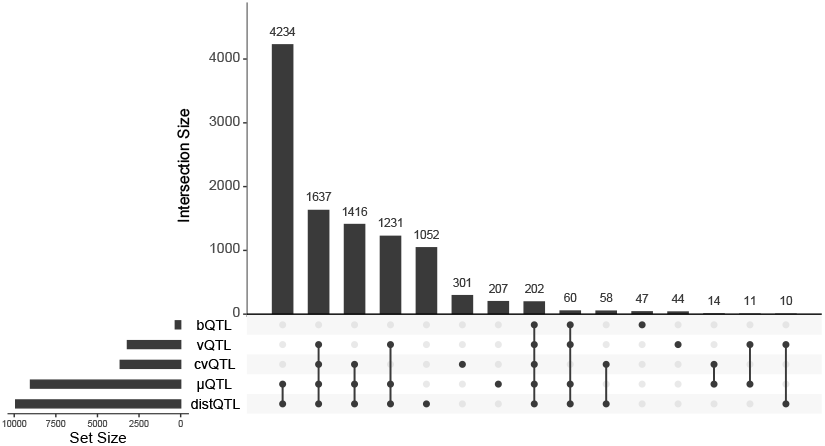
Upset plot of eQTL-labeled gene-SNP pairs across B cells, using corrected p-values and significance threshold *α* = 10^−4^. Truncated to 15 intersections.

#### distQTL v.s. PMER

We compare distQTL to PMER (6) on the top eQTLs identified by distQTL in B cells. Fig. 5 illustrates log_10_(− log_10_ *p*) transformed nominal p-values from distQTL and PMER, using only highly expressed genes in a cell-type-specific manner. The transformation visually separates significant p-values while reducing skew: large positive values indicate highly significant test results, e.g. a value of 1 corresponds to *p* = 1E−10, and a value of 0 corresponds to *p* = 0.1. In total, we fit 35,035 cell-type-specific PMER models, using the Texas A&M High Performance Research Computing (TAMU-HPRC) resource.

**Fig. 5.**
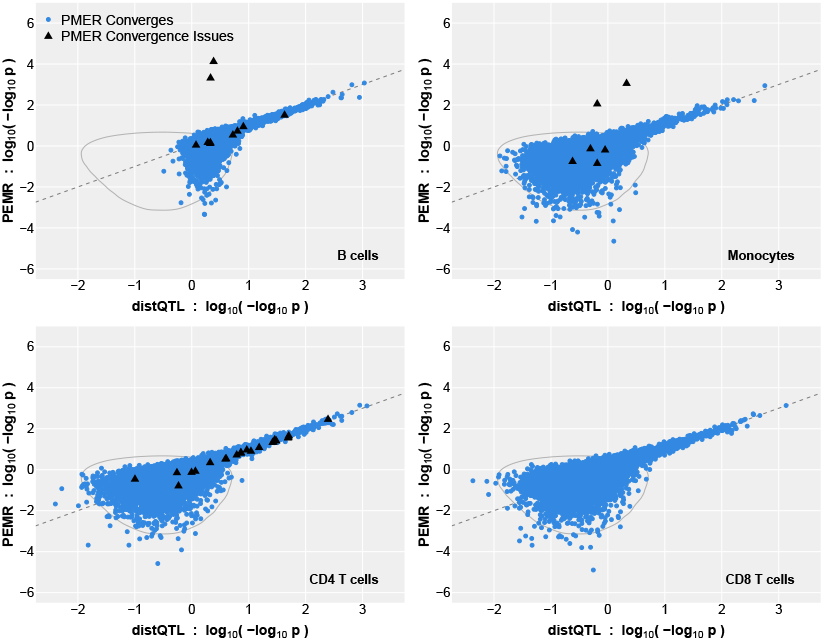
eQTL-mapping log_10_ (− log_10_ *p*)-value comparison between PMER and distQTL for each cell-type group. Each point corresponds to one highly expressed gene and its top eQTL identified by distQTL. Contours are visual aids for joint p-value distribution under null of no SNP effect. PMER fits accompanied by convergence flags are labeled with black triangles. For reference, PMER model fit time was approximately 33 CPU hours; distQTL model fit time was approximately 1.6 CPU minutes.

Despite our selection of top eQTLs using distQTL biasing significance toward distQTL, distQTL and PMER visually exhibit strong correlation in eQTL-mapping results and attain very similar p-values when identifying significant eQTLs. However, on occasion the PMER model fitting procedure does not converge. The most common convergence issues are violation of the stationarity condition, and non-convexity or near-singularity of the Hessian, indicating poor identification of some parameters. These convergence issues occurred despite scaling covariate vectors prior to model fit. PMER convergence problems do not necessarily mean poor estimated coefficients, but sometimes produce unrealistically significant eQTL identification. Spurious results are only identifiable by comparison to other methods, or otherwise by resolving the convergence problems through manual tuning of the optimization algorithm on a case-by-case basis. While both methods are visually comparable on significance calls in Fig. 5, the convergence issues and computational bottleneck of PMER make distQTL a more appealing standalone scRNA-seq method.

### Benchmarking and Validation

Baseline eQTL methods and distQTL applied to OneK1K data set for cell-type group exhibit varying levels of enrichment against eQTL findings from whole blood bulk tissue samples in GTEx. We compare methods individually to GTEx findings by identifying *cis*-SNPs as eQTLs if the corresponding eQTL-mapping method’s corrected (as applicable) p-value is smaller than 10^−4^, and counting matches of eQTL labeling between the two data sets. To reduce noise for illustrative purposes, we narrow focus to genes with at least four eQTLs found in both GTEx and OneK1K, on a method-by-method basis.

Fig. 6**A** shows contour plots of within-gene proportion of OneK1K eQTL labels which are also labeled eQTLs in GTEx, against proportion of GTEx gene-SNP pairs which were eQTL labeled. Values above the diagonal line indicate better than chance matching of GTEx-labeled eQTLs by methods applied to the OneK1K data set. Baseline *µ*QTL and distQTL outperform the other methods both in identifying eQTLs in the aggregated cell groups and in identifying eQTLs that are also eQTLs in GTEx. Across all gene-SNP pairs that were identified as significant by GTEx, *µ*QTL in OneK1K recovered 24.1%, and distQTL recovered 23.1%. We note that aggregate replication is likely hampered by mismatching blood sample characteristics, as whole blood samples from GTEx contain granulocyte and other cell types which are not present in the PBMC-only samples from the OneK1K cohort (7; 32). When using cell-type-specific results, cell types with higher relative abundance in blood samples exhibit higher eQTL-mapping overlap with GTEx; see Supplement (Table S3).

**Fig. 6.**
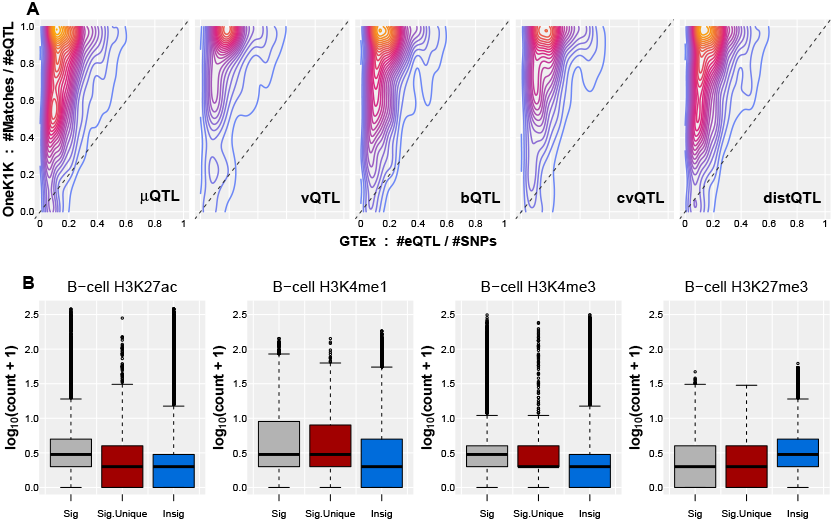
Benchmark and validation results. (**A**) Gene-SNP OneK1K/GTEx enrichment contour plots, conditioned on OneK1K eQTL ≥ 4 and GTEx eQTL ≥ 4. eQTLs identified by *p* ≤ 10^−4^; matches are eQTL identification in both data sets. Values above the dashed *x* = *y* line indicate eQTL matching better than chance. (**B**) B-cell eQTL validation by ChIP-seq of histone modifications. Sig.: significant QLTs detected by distQTL. Sig.Unique: Significant eQTLs detected exclusively by distQTL (not identified by other methods). Insig.: insignificant eQTLs. log_10_(*y* + 1) of ChIP-seq read counts is shown as y-axis. eQTLs detected (uniquely) by distQTL show stronger ChIP-seq signals for enhancer markers (H3K27ac, H3K4me1, and H3K4me3) and weaker signals for repression marker (H3K27me3), demonstrating the accuracy and outperformance of distQTL.

To further benchmark distQTL against the other univariate methods, we adopted cell-type-specific epigenomic profiles, encompassing ChIP-seq data of histone modifications. Compared to non-significant calls, eQTLs detected by distQTL overall and eQTLs exclusively detected by distQTL (but not by the other methods) exemplify stronger signals for active enhancer markers and weaker signal for repression marker (Fig. 6**B**). This orthogonal high-throughput validation from a different modality demonstrates the accuracy and outperformance of distQTL.

### Computation Time

Table 1 displays model fit times and total number of models fit, in each cell-type group, by competing eQTL-mapping methods. Pseudo-bulk methods reduce scRNA-seq data set size by collapsing donor-level gene expression data into single numeric summaries. For simplicity, we present results only for *µ*QTL; computation results for other baseline eQTL-mapping methods are nearly identical. Estimated time to fit each model with *µ*QTL is on the order of 0.7 ms, owing to closed form solutions. Total *µ*QTL run time across 5.8 million models was approximately 8 minutes.

**Table 1.** Model fit times (in minutes) for high expression genes, within each cell-type group; total number of models fit per method across the four cell-type groups; and average model fit time (in milliseconds).

| Method | Total 🕒 (min) |  |  |  | Num. Models | Ave. 🕒 (ms) |
| --- | --- | --- | --- | --- | --- | --- |
|  | B | Mono | CD4 T | CD8 T |  |  |
| μQTL | 2.1 | 1.6 | 2.1 | 2.1 | 5.8 M | 7.3E-1 |
| PMER | 43.7 | 18.3 | 90.3 | 60.3 | 4000 | 2.9E+4 |
| distQTL | 65.3 | 101.6 | 37.5 | 60.7 | 5.8 M | 2.5E+1 |

scRNA-seq specific methods try to leverage the complete data set, which can increase computational burden. For PMER method, each individual cell is present as a row in the response and covariate objects, with a random intercept per donor accounting for cell-cell correlations. Such model structure requires very large data object sizes, with a large number of non-trivial parameters to estimate per model. To comparably evaluate PMER to other methods on the same machine, we fit a reduced set of PMER models as baseline eQTL and distQTL, using parallel computing. We randomly selected 1000 genes which have high expression in all cell-type groups, and fit the top eQTL per gene as identified by distQTL in B cells. Average model fit time was approximately 29 seconds, upward of one minute for cell-type groups with larger numbers of cells. Total time to fit all 4000 PMER models was 33 CPU hours. (Fitting the larger set of 35,035 models on TAMU-HPRC took approximately 589 CPU hours.) Notably, model fit time was weakly negatively correlated with eQTL significance level, indicating our reported average model fit times are likely optimistic lower bounds if PMER is applied to all possible pairs of genes and *cis*-SNPs in OneK1K, some 2.9 million total models. This presents a daunting computational task, even with access to high power computational resources like TAMU-HPRC, and is infeasible with a typical personal computer.

distQTL is an scRNA-seq method that replaces complicated covariance structures with donor-specific distribution responses. Performing the partial *F* -test involves matrix calculations that have closed form solutions, provided the conditional mean distributions from eq. (5) are calculated. The bulk of distQTL computational burden correspondingly lies in solving eq. (5), a convex quadratic programming problem (26) which can be quickly solved to machine precision (33). Individual distQTL model fit times are on the order of tens of milliseconds, upward of 1000*×* reduction in computation time compared to PMER, and obtaining more numerically stable results. On a personal laptop, total time to fit 5.8 million high expression gene-SNP models, i.e. including with permuted responses, was a little over four hours. We believe this makes distQTL an attractive and computationally feasible scRNA-seq method which sidesteps the cell-cell correlation issue at the donor level.

## Discussion

Here, we propose distQTL for eQTL-mapping using population-scale scRNA-seq data, demonstrating its high power to detect mean effects while also identifying non-mean associations. Simulations (Fig. 1) and application to the OneK1K data set (Table S1; Table S2) suggest that distQTL can identify more eQTLs that other methods lack the power to find. Aggregating p-value results from permutation-refitted models indicates that type-I errors remain well-controlled. As with other eQTL-mapping methods, refitting models under permutation is a computational bottleneck, and the method cannot adjust very significant p-values. Visual inspection of the uncorrected p-value distribution in Fig. 3**B** suggests that beta distribution correction methods (34) will not be appropriate. Nonetheless, fast implementation of Fréchet regression enables efficient model fitting under permutation, reducing computation time to just a few hours on a laptop. In OneK1K, we have over 700,000 gene-SNP pairs per cell-type group; applying a single permutation p-value per model allows us to use pooled correction (29) for p-values far below our stringent significance threshold of *p* ≤ 10^−4^. Further, while dependent p-values from pseudo-bulk approaches (e.g., *µ*QTL, vQTL, bQTL, and cvQTL) can be integrated using heavy-tailed combination tests (35), it is not clear the pooling correction method, applied independently to each method’s p-values, is sufficient for joint correction prior to combination test usage. This is a benefit of distQTL as a unified method that extends beyond the first and second moments of the gene expression distribution.

Tissue-specific eQTL-mapping by the GTEx Project includes more sophisticated regressors such as the PEER factors (2), which have been shown to increase power of eQTL-mapping by modeling latent batch effects (36). A recent paper (37) suggests gene expression PCA is another, faster latent correction method comparable to PEER factors for eQTL-mapping. However, we do not include specific latent batch effect correction in our exposition of distQTL due to complications in applying these methods to population-scale scRNA-seq data that exhibit nested structure. Choosing QC steps and optimal number of latent factors with averaged pseudo-bulk samples has been shown to be non-trivial (38) with heterogeneous results across cell-types, whether by PEER factors or gene expression PCA. Additionally, at the time of writing we were not able to install the R peer package which implements the PEER factor analysis through the Comprehensive R Archive Network or its GitHub repository. Nonetheless, we emphasize that while batch effect correction is a nuanced problem that we do not tackle, distQTL is amenable to inclusion of such other donor-level covariates. Despite utilizing a simpler regression model across methods, distQTL identifies more gene-SNP pairs in OneK1K which are significant at the 10^−4^ level while *µ*QTL (and occasionally the other univariate methods) retains non-significance at the 0.10 level (Table S2).

For scRNA-seq data, transcriptional bursting that results in stochastic gene expression is a widespread phenomenon observed at the single-cell level, and a two-state Poisson-Beta model has been successfully adopted to infer bursting kinetics (21; 39). However, data from 5’ or 3’ end tagging techniques (e.g., 10x Genomics) is sparser and noisier than full-transcript scRNA-seq, leading to poor model fittings and identifiability issues (21; 40). The inherent heterogeneity across cell types and/or cell states further exacerbates the problem, necessitating a large number of cells for accurate parameter estimation. While we included and benchmarked bursting QTLs – using the proportion of cells with non-zero expression as a surrogate of the bursting frequency, which cannot be uniquely and accurately estimated – distQTL characterizes conditional gene distributions beyond mere univariate reductions or parametric forms. Our method does not make parametric assumptions on the gene expression and hence avoids the need to separately model zeros in a hurdle model (41; 42), implement zero-inflation through parametric assumptions (21; 39), or rely on zero-imputation methods (43; 44; 45; 46; 47) whose uncertainty is non-trivial to account for in downstream inference. Additionally, and perhaps more importantly for the OneK1K cohort, separate zero-inflation modeling for technical zeros has been shown to be unnecessary for UMI-based scRNA-seq data (48; 49), and observed counts appear sufficient for many inference tasks (50). This makes distQTL an appealing low-assumption, omnibus approach that can flexibly handle multimodal or sparse gene expression data sets.

Rapidly increasing sequencing capacity and technological advancement implies distQTL – alongside univariate summaries such as variance, bursting kinetics, and coefficient of variation when applied to more cells sequenced at a higher depth – can uncover regulatory relationships at a finer resolution. distQTL is not limited to using expression as the quantitative trait; it can be readily extended to other omics measures such as chromatin accessibility, alternative splicing, and allele-specific expression. Mapping eQTLs using single-cell multiomics data, where multiple modalities are simultaneously profiled within the same cells (51), holds the potential to provide deeper insights into into the intricate transcriptional regulatory network and warrants additional methodological developments. Marginal effects on responses – including the univariate summaries we consider (i.e., mean, variance, positive proportion, or measures of dispersion) – can be immediately obtained from conditional mean distributions obtained from distQTL. While distQTL testing is an omnibus method, methods for utilizing distributional summaries for biological mechanism recovery warrant further investigation.

## Data availability

OneK1K scRNA-seq and genotype data were downloaded from the HCA (https://explore.data.humancellatlas.org/projects/f2078d5f-2e7d-4844-8552-f7c41a231e52). H3K27ac, H3K4me1, H3K4me3, and H3K27me3 ChIP-seq data of B cells (CD19) were downloaded from GEO with accession numbers GSM1027287, GSM1027296, GSM1027300, and GSM1160194, respectively. Tissue-specific *cis*-QLTs from GTEx Analysis V8 were downloaded from the GTEx Portal (https://www.gtexportal.org/home/downloads/adult-gtex/qtl).

## Competing interests

The authors declare no competing interests.

## Author contributions statement

A.C., Y.N., and Y.J. initiated the study and conceived the methods. A.C. and Y.J. performed data quality control; all authors performed data analysis and results interpretation. Y.N. and Y.J. supervised the work and provided funding support. A.C. and Y.J. wrote the manuscript, which was reviewed and approved by all authors. A.C. wrote the accompanying R package; a development version is available at https://github.com/alexandercoulter/distQTL.

## Acknowledgments

This work was supported by the National Institute of Health (NIH) grant R01 GM148974 (to Y.N.) and R35 GM138342 (to Y.J.). The authors thank Dr. Joseph Powell for providing support on the OneK1K genotype calls. Portions of this research were conducted with the advanced computing resources and consultation provided by Texas A&M High Performance Research Computing.

## Notes

https://explore.data.humancellatlas.org/projects/f2078d5f-2e7d-4844-8552-f7c41a231e52

### Competing Interest Statement

The authors have declared no competing interest.

### Summary of Updates

Updated literature review, simulation settings, and discussion of results.

## References

1. Mackay, T. F. C., Stone, E. A., and Ayroles, J. F. The genetics of quantitative traits: challenges and prospects. Nat. Rev. Genet. 2009; 10(8): 565–577.

2. THE GTEX CONSORTIUM The GTEx Consortium atlas of genetic regulatory effects across human tissues. Science 2020; 369(6509): 1318–1330.

3. van der Wijst, M. G. P., Brugge, H., de Vries, D. H., et al. Single-cell RNA sequencing identifies celltype-specific cis-eQTLs and co-expression QTLs. Nat. Genet. 2018; 50(4): 493–497.

4. Neavin, D., Nguyen, Q., Daniszewski, M. S., et al. (2021) Single cell eQTL analysis identifies cell type-specific genetic control of gene expression in fibroblasts and reprogrammed induced pluripotent stem cells. Genome Biol., 22(1), 76. 10.1186/s13059-021-02293-3

5. Jerber, J., Seaton, D. D., Cuomo, A. S. E., et al. Population-scale single-cell RNA-seq profiling across dopaminergic neuron differentiation. Nat. Genet. 2021; 53(3): 304–312.

6. Nathan, A., Asgari, S., Ishigaki, K., et al. Single-cell eQTL models reveal dynamic T cell state dependence of disease loci. Nature 2022; 606(7912): 120–128.

7. Yazar, S., Alquicira-Hernandez, J., Wing, K., et al. (2022) Single-cell eQTL mapping identifies cell type–specific genetic control of autoimmune disease. Science, 376(6589), eabf3041. 10.1126/science.abf3041

8. Maria, M., Pouyanfar, N., Ö rd, T., et al. (2022) The Power of Single-Cell RNA Sequencing in eQTL Discovery. Genes, 13(3), 502. 10.3390/genes13030502

9. Luo, J., Wu, X., Cheng, Y., et al. (2023) Expression quantitative trait locus studies in the era of single-cell omics. Front. Genet., 14, 1182579. 10.3389/fgene.2023.1182579

10. Ding, R., Wang, Q., Gong, L., et al. (2024) scQTLbase: an integrated human single-cell eQTL database. Nucleic Acids Res., 52(D1), D1010–D1017. 10.1093/nar/gkad781

11. Elorbany, R., Popp, J. M., Rhodes, K., et al. (2022) Single-cell sequencing reveals lineage-specific dynamic genetic regulation of gene expression during human cardiomyocyte differentiation. PLoS Genet., 18(1), e1009666. 10.1371/journal.pgen.1009666

12. Corty, R. W. and Valdar, W. vqtl: An R Package for Mean-Variance QTL Mapping. G3 Genes|Genomes|Genetics 2018; 8(12): 3757–3766.

13. Hong, C., Ning, Y., Wei, P., et al. A Semiparametric Model for VQTL Mapping. Biometrics 2017; 73(2): 571–581.

14. Sarkar, A. K., Tung, P.-Y., Blischak, J. D., et al. (2019) Discovery and characterization of variance QTLs in human induced pluripotent stem cells. PLoS Genet., 15(4), e1008045. 10.1371/journal.pgen.1008045

15. Eling, N., Richard, A. C., Richardson, S., et al. Correcting the Mean-Variance Dependency for Differential Variability Testing Using Single-Cell RNA Sequencing Data. Cell Syst. 2018; 7(3): 284–294.e12.

16. Westerman, K. E., Majarian, T. D., Giulianini, F., et al. (2022) Variance-quantitative trait loci enable systematic discovery of gene-environment interactions for cardiometabolic serum biomarkers. Nat. Commun., 13(1), 3993. 10.1038/s41467-022-31625-5

17. Shi, G. (2022) Genome-wide variance quantitative trait locus analysis suggests small interaction effects in blood pressure traits. Sci. Rep., 12(1), 12649. 10.1038/s41598-022-16908-7

18. Rönnegård, L. and Valdar, W. Detecting Major Genetic Loci Controlling Phenotypic Variability in Experimental Crosses. Genetics 2011; 188(2): 435–447.

19. Forsberg, S. K. G., Andreatta, M. E., Huang, X.-Y., et al. (2015) The Multi-allelic Genetic Architecture of a Variance-Heterogeneity Locus for Molybdenum Concentration in Leaves Acts as a Source of Unexplained Additive Genetic Variance. PLoS Genet., 11(11), e1005648. 10.1371/journal.pgen.1005648

20. Larsson, A. J. M., Johnsson, P., Hagemann-Jensen, M., et al. Genomic encoding of transcriptional burst kinetics. Nature 2019; 565(7738): 251–254.

21. Jiang, Y., Zhang, N. R., and Li, M. (2017) SCALE: modeling allele-specific gene expression by single-cell RNA sequencing. Genome Biol., 18(1), 74. 10.1186/s13059-017-1200-8

22. Li, S., Guo, D., Sun, Q., et al. MAPK4 silencing in gastric cancer drives liver metastasis by positive feedback between cancer cells and macrophages. Exp. Mol. Med. 2023; 55(2): 457–469.

23. Jia, C., Wang, L. Y., Yin, G. G., et al. (2019) Single-cell stochastic gene expression kinetics with coupled positiveplus-negative feedback. Phys. Rev. E, 100(5), 052406. 10.1103/PhysRevE.100.052406

24. Chuffart, F., Richard, M., Jost, D., et al. (2016) Exploiting Single-Cell Quantitative Data to Map Genetic Variants Having Probabilistic Effects. PLoS Genet., 12(8), e1006213. 10.1371/journal.pgen.1006213

25. Zhang, M., Liu, S., Miao, Z., et al. (2022) IDEAS: individual level differential expression analysis for singlecell RNA-seq data. Genome Biol., 23(1), 33. 10.1186/s13059-022-02605-1

26. Petersen, A. and Müller, H.-G. Fréchet regression for random objects with Euclidean predictors. Ann. Stat. 2019; 47(2): 691–719.

27. Shen, Q. and Faraway, J. An F Test for Linear Models with Functional Responses. Stat. Sin. 2004; 14(4): 1239–1257.

28. Petersen, A., Liu, X., and Divani, A. A. Wasserstein F-tests and confidence bands for the Fréchet regression of density response curves. Ann. Stat. 2021; 49(1): 590–611.

29. Storey, J. D. and Tibshirani, R. Statistical significance for genomewide studies. Proc. Natl. Acad. Sci. 2003; 100(16): 9440–9445.

30. Bates, D., Mächler, M., Bolker, B., et al. (2015) Fitting Linear Mixed-Effects Models Using lme4. J. Stat. Software, 67(1), 1–48. 10.18637/jss.v067.i01

31. Bernstein, B. E., Stamatoyannopoulos, J. A., Costello, J. F., et al. The NIH Roadmap Epigenomics Mapping Consortium. Nat. Biotechnol. 2010; 28(10): 1045–1048.

32. Pagana, K. D., Pagana, T. J., and Pagana, T. N. Mosby’s® diagnostic and laboratory test reference. St. Louis, MO: Mosby, 2024, 17 edition.

33. Coulter, A., Lee, R., and Gaynanova, I. (2025) fastfrechet: An R package for fast implementation of Fréchet regression with distributional responses. J. Open Source Software, 10(109), 7925. 10.21105/joss.07925

34. Ongen, H., Buil, A., Brown, A. A., et al. Fast and efficient QTL mapper for thousands of molecular phenotypes. Bioinformatics 2016; 32(10): 1479–1485.

35. Gui, L., Jiang, Y., and Wang, J. (2025) Aggregating Dependent Signals with Heavy-Tailed Combination Tests. Biometrika, p. asaf038 10.1093/biomet/asaf038

36. Stegle, O., Parts, L., Durbin, R., et al. (2010) A Bayesian Framework to Account for Complex Non-Genetic Factors in Gene Expression Levels Greatly Increases Power in eQTL Studies. PLoS Comput. Biol., 6(5), e1000770. 10.1371/journal.pcbi.1000770

37. Zhou, H. J., Li, L., Li, Y., et al. (2022) PCA outperforms popular hidden variable inference methods for molecular QTL mapping. Genome Biol., 23(1), 210. 10.1186/s13059-022-02761-4

38. Xue, A., Yazar, S., Neavin, D., et al. (2023) Pitfalls and opportunities for applying latent variables in singlecell eQTL analyses. Genome Biol., 24(1), 33. 10.1186/s13059-023-02873-5

39. Vu, T. N., Wills, Q. F., Kalari, K. R., et al. Beta-Poisson model for single-cell RNA-seq data analyses. Bioinformatics 2016; 32(14): 2128–2135.

40. Weideman, A. M. K., Wang, R., Ibrahim, J. G., et al. (2025) Canopy2: Tumor Phylogeny Inference by Bulk DNA and Single-Cell RNA Sequencing. Stat. Biosci., 10.1007/s12561-024-09466-1

41. Finak, G., McDavid, A., Yajima, M., et al. (2015) MAST: a flexible statistical framework for assessing transcriptional changes and characterizing heterogeneity in single-cell RNA sequencing data. Genome Biol., 16(1), 278. 10.1186/s13059-015-0844-5

42. Wang, J., Huang, M., Torre, E., et al. Gene expression distribution deconvolution in single-cell RNA sequencing. Proc. Natl. Acad. Sci. 2018; 115(28): E6437–E6446.

43. Eraslan, G., Simon, L. M., Mircea, M., et al. (2019) Singlecell RNA-seq denoising using a deep count autoencoder. Nat. Commun., 10(1), 390. 10.1038/s41467-018-07931-2

44. Chen, M. and Zhou, X. (2018) VIPER: variability-preserving imputation for accurate gene expression recovery in single-cell RNA sequencing studies. Genome Biol., 19(1), 196. 10.1186/s13059-018-1575-1

45. van Dijk, D., Sharma, R., Nainys, J., et al. Recovering Gene Interactions from Single-Cell Data Using Data Diffusion. Cell 2018; 174(3): 716–729.e27.

46. Huang, M., Wang, J., Torre, E., et al. SAVER: gene expression recovery for single-cell RNA sequencing. Nat. Methods 2018; 15(7): 539–542.

47. Li, W. V. and Li, J. J. (2018) An accurate and robust imputation method scImpute for single-cell RNA-seq data. Nat. Commun., 9(1), 997. 10.1038/s41467-018-03405-7

48. Svensson, V. Droplet scRNA-seq is not zero-inflated. Nat. Biotechnol. 2020; 38(2): 147–150.

49. Sarkar, A. and Stephens, M. Separating measurement and expression models clarifies confusion in single-cell RNA sequencing analysis. Nat. Genet. 2021; 53(6): 770–777.

50. Jiang, R., Sun, T., Song, D., et al. (2022) Statistics or biology: the zero-inflation controversy about scRNA-seq data. Genome Biol., 23(1), 31. 10.1186/s13059-022-02601-5

51. Jiang, Y., Harigaya, Y., Zhang, Z., et al. Nonparametric single-cell multiomic characterization of trio relationships between transcription factors, target genes, and cisregulatory regions. Cell Syst. 2022; 13(9): 737–751.e4.

