## Supplement for "distQTL: Distribution Quantitative Trait Loci Identification by Population-Scale Single-Cell Data"

### S1 distQTL Partial F-tests

#### S1.1 Variant from Petersen, Liu, and Divani (2021)

Let $\mathcal{Q}$ be the space of univariate quantile functions $\{Q\}$ equipped with the 2-Wasserstein metric $d_W(\mathbf{p},\mathbf{q}) := \|\mathbf{p}-\mathbf{q}\|_{L^2}$, and let $X = (Y^\top \quad Z^\top)^\top \in \mathbb{R}^{p_y+p_z}$ be a covariate vector with $\mathbb{E}(X) = \mathbf{0}$ and variance-covariance matrix

$$\mathrm{Var}(X) \equiv \mathbf{\Sigma}_X \equiv \begin{pmatrix} \mathbf{\Sigma}_Y & \mathbf{\Sigma}_{YZ} \\ \mathbf{\Sigma}_{ZY} & \mathbf{\Sigma}_Z \end{pmatrix}.$$

Let $p := p_y + p_z$. Assume we observe sample pairs $\{\mathbf{x}_i, Q_i\}_{i=1}^n$, where covariate matrix $\mathbf{X} := (\mathbf{x}_1 \quad \cdots \quad \mathbf{x}_n)^\top \in \mathbb{R}^{n\times p}$ has been column-centered. To test the null hypothesis $H_0 : Q_\oplus(X) = Q_\oplus(Y)$ (i.e. accounting for $Y$, component $Z$ has no association with the conditional Fréchet mean), [1] propose a partial $F$-test statistic

$$F := \sum_{i=1}^n \left[ d_W^2 \left\{ \widehat{Q}_\oplus(\mathbf{x}_i), \widehat{Q}_\oplus(\overline{\mathbf{x}}) \right\} - d_W^2 \left\{ \widehat{Q}_\oplus(\mathbf{y}_i), \widehat{Q}_\oplus(\overline{\mathbf{x}}) \right\} \right], \tag{S1}$$

where $\widehat{Q}_\oplus(\mathbf{x}_i)$ is the solution to the $i^{\text{th}}$ Fréchet regression problem for the full model, $\widehat{Q}_\oplus(\mathbf{y}_i)$ is the solution for the reduced model, and $\widehat{Q}_\oplus(\overline{\mathbf{x}})$ is the solution for the "intercept-only" model. Letting

$$\mathbf{J}^\top := (-\mathbf{\Sigma}_{ZY}\mathbf{\Sigma}_Y \mid \mathbf{I}_{p_z}), \qquad \mathbf{\Sigma}_{Z|Y} := \mathbf{\Sigma}_Z - \mathbf{\Sigma}_{ZY}\mathbf{\Sigma}_Y^{-1}\mathbf{\Sigma}_{YZ},$$

[1] define covariance kernel

$$C(u,v) := \mathbf{\Sigma}_{Z|Y}^{-1/2}\mathbf{J}^\top \mathbb{E}\left[ XX^\top \mathrm{Cov}\left\{Q(X;u), Q(X;v)\right\}\right] \mathbf{J}\mathbf{\Sigma}_{Z|Y}^{-1/2}, \qquad u,v \in (0,1).$$

They show that under certain regularity conditions (namely, absolute continuity of the measures associated with $Q(X)$, and conditions on $\mathbb{E}(Z|Y)$ and $\mathrm{Var}(Z|Y)$), the asymptotic distribution of $F$ can be characterized

$$F|\{X_i\}_{i=1}^n \xrightarrow{\mathcal{D}} \sum_{j=1}^\infty \lambda_j \xi_j^2,$$

where the $\lambda_j$'s are the eigenvalues of kernel $C$, and $\xi_j \overset{\text{iid}}{\sim} \mathcal{N}(0,1)$. [1] approximate the null distribution of $F$ with a scaled $\chi^2$ distribution $a\chi_b^2$ [2], where $a, b$ satisfy

$$ab = p\sum_{j=1}^\infty \lambda_j = p\int_0^1 C(u,u)du, \qquad a^2 b = p\sum_{j=1}^\infty \lambda_j^2 = p\int_0^1\int_0^1 C(u,v)dudv.$$

$C$ is estimated with plug-in values $\widehat{\mathbf{\Sigma}}_{Z|Y}$ and $\widehat{\mathbf{J}}$, and typically quantile functions are evaluated on a common $m$-grid in $(0,1)$, i.e. $u, v \in \{\frac{1}{2m}, \frac{3}{2m}, \cdots, 1-\frac{1}{2m}\}$. The final estimator [1] use is

$$\widehat{C}(u,v) := \widehat{\mathbf{\Sigma}}_{Z|Y}^{-1/2}\widehat{\mathbf{J}}^\top \left[ \frac{1}{n}\sum_{i=1}^n \mathbf{x}_i\mathbf{x}_i^\top \left\{ Q_i(u) - \widehat{Q}_\oplus(\mathbf{x};u)\right\}\left\{ Q_i(v) - \widehat{Q}_\oplus(\mathbf{x};v)\right\}\right] \widehat{\mathbf{J}}\widehat{\mathbf{\Sigma}}_{Z|Y}^{-1/2}, \tag{S2}$$

where we denote $\widehat{\mathbf{C}}$ as the matrix whose $(u, v)$ entry is (S2). The values of $\widehat{a}, \widehat{b}$ solving the system above for $\widehat{\mathbf{C}}$ can be found by

$$\widehat{a} = \frac{\frac{1}{m^2}\|\widehat{\mathbf{C}}\|_F^2}{\frac{1}{m}\mathrm{tr}(\widehat{\mathbf{C}})}, \qquad \widehat{b} = \frac{1}{\widehat{a}} \times \frac{1}{m}\mathrm{tr}(\widehat{\mathbf{C}}).$$

The $\alpha$-level test for $H_0$ is thus to reject $H_0$ *iff*

$$F \geq \widehat{a} \times Q_{\chi^2_{\widehat{b}}}(1-\alpha) \qquad \Longleftrightarrow \qquad P_F := 1 - F_{\chi^2_{\widehat{b}}}(F/\widehat{a}) \leq \alpha, \tag{S3}$$

where $Q_{\chi^2_{\widehat{b}}}$ and $F_{\chi^2_{\widehat{b}}}$ are respectively the quantile function and CDF of the $\chi^2$ distribution with degrees of freedom $\widehat{b}$, and $P_F$ is the p-value.

#### S1.2 Variant from Shen and Faraway (2004)

The $F$ statistic (S1) of [1] is analogous to the numerator of the partial $F$-test statistic in classical linear regression, and correspondingly their implementation of the Satterthwaite approximation of [2] is for this quasi-numerator only. The proposed statistic in [2] includes a denominator explicitly, with its own Satterthwaite approximation. In the context of Fréchet regression using sample covariate-response pairs $\{(\mathbf{x}_i, Q_i)\} \subset \mathbb{R}^p \times \mathcal{Q}$, the proposed test statistic from [2] has the form

$$F_S := \frac{n-p}{p_z} \times \frac{R_0 - R_1}{R_1},$$

where

$$R_0 := \sum_{i=1}^n d_W^2 \left\{\widehat{Q}_\oplus(\mathbf{y}_i), Q_i\right\}, \qquad R_1 := \sum_{i=1}^n d_W^2 \left\{\widehat{Q}_\oplus(\mathbf{x}_i), Q_i\right\}.$$

The limiting random variable under $H_0$ is then approximated with

$$\lim_{n\to\infty} F_S|\{X_i\}_{i=1}^n \approx \frac{n-p}{p_z} \times \frac{a_1\phi_1}{a_2\phi_2}$$

where $\phi_1 \sim \chi^2_{b_1}$ and $\phi_2 \sim \chi^2_{b_2}$; the parameters $a_1, b_1$ and $a_2, b_2$ satisfy

$$\frac{a_1}{p_z}b_1 = p\int_0^1 C(u,u)du, \qquad \frac{a_1^2}{p_z^2}b_1 = p\int_0^1\int_0^1 C(u,v)dudv,$$
$$\frac{a_2}{n-p}b_2 = p\int_0^1 C(u,u)du, \qquad \frac{a_2^2}{(n-p)^2}b_2 = p\int_0^1\int_0^1 C(u,v)dudv;$$

and $C(u, v)$ is estimated with (S2). Note in the sample setting $\widehat{a}_1 = \widehat{a}_2$, and

$$\widehat{b}_1 = p_z\frac{\mathrm{tr}(\widehat{\mathbf{C}})^2}{\|\widehat{\mathbf{C}}\|_F^2}, \qquad \widehat{b}_2 = (n-p)\frac{\mathrm{tr}(\widehat{\mathbf{C}})^2}{\|\widehat{\mathbf{C}}\|_F^2}.$$

We can see $\widehat{b}_2/\widehat{b}_1 = (n-p)/p_z$, hence

$$\frac{n-p}{p_z} \times \frac{\widehat{a}_1\phi_1}{\widehat{a}_2\phi_2} = \frac{\phi_1/\widehat{b}_1}{\phi_2/\widehat{b}_2} \sim \mathrm{F}_{\widehat{b}_1,\widehat{b}_2}.$$

The $\alpha$-level test for $H_0$ is thus to reject $H_0$ *iff*

$$F_S \geq Q_{\mathrm{F}_{\widehat{b}_1,\widehat{b}_2}}(1-\alpha) \qquad \Longleftrightarrow \qquad P_{F_S} := 1 - F_{\mathrm{F}_{\widehat{b}_1,\widehat{b}_2}}(F_S) \leq \alpha, \tag{S4}$$

where $Q_{\mathrm{F}_{\widehat{b}_1,\widehat{b}_2}}$ and $F_{\mathrm{F}_{\widehat{b}_1,\widehat{b}_2}}$ are respectively the quantile function and CDF of the F distribution with degrees of freedom parameters $\widehat{b}_1$ and $\widehat{b}_2$.

#### S1.3 Performance Testing Effects of Very Rare Alleles

In the context where we are testing for a SNP effect on gene expression while controlling for other covariates, our observed covariate vectors take form $\mathbf{x}_i = (\mathbf{y}^\top \quad z)^\top$, where $z \in \{0, 1, 2\}$ corresponds to SNP genotype $AA$, $Aa$, and $aa$, respectively. A minor allele can be very rare, i.e. most observations are $z = 0$, with only a handful of $z = 1, 2$ observations. The asymptotic null distributions of test statistics like $F$ and $F_S$ will not necessarily be appropriate in these settings, owing to noisy estimation of covariance objects like $\mathbf{\Sigma}_{Z|Y}$.

We evaluate the finite sample performance of $F$ and $F_S$ under the null hypothesis $H_0 : Q_\oplus(\mathbf{x}) = Q_\oplus(\mathbf{y})$ by generating $n = 1000$ random zero-inflated negative binomial (`zinbinom`) distributions, represented by discretized quantile functions $\{Q_i\}$. For each $Q_i$, we sample covariate vector $\mathbf{x}_i = (\mathbf{y}_i^\top \quad z_i)^\top \in \mathbb{R}^5$, where $\mathbf{y}_i \overset{\text{iid}}{\sim} \mathcal{N}(\mathbf{0}, \mathbf{I}_4)$, and independently we sample "SNP" genotype $z_i = 1$ (zero otherwise) for randomly selected index subset $\{i\} \equiv \mathcal{I} \subset \{1, \ldots, n\}$; by design, $Q_i$ is not dependent on $\mathbf{x}_i$. We repeat this simulation process 10,000 times for each $|\mathcal{I}| \in \{1, 2, \ldots, 10\}$, and perform statistical tests (S3) and (S4) for the SNP effect.

Figure S1 compares QQ-plots of p-values from the two testing methods in the presence of increasing incidence of "rare allele" observations, i.e. $|\mathcal{I}| = 1, 2, \ldots, 10$, against the expected quantiles from the typical unif$(0, 1)$ distribution under a correctly specified null distribution. While neither method gives a uniform p-value distribution, since the `zinbinom` distribution is not absolutely continuous (with respect to the Lebesgue measure), the testing method from [1] using the $F$ statistic has greater finite-sample sensitivity to $|\mathcal{I}|$. In contrast, the p-value distribution using the testing method from [2] exhibits stability across small values of $|\mathcal{I}|$. Both methods have the greatest distortion at the edge case $|\mathcal{I}| = 1$.

For this manuscript, we utilize the testing procedure (S4) of [2], and restrict eQTL finding to those gene-SNP pairs for which the rare SNP allele is present in at least 5 donors (out of 961 donors at most). This reduces the finite-sample dependence of distQTL on the covariate matrix $\mathbf{X}$.

#### S1.4 Illustrating Linear and Distributional F-tests

We illustrate the basic idea of distributional regression by comparison to linear regression for a two-covariate toy example in Figure S2. For this toy example, the covariate of interest $X_2$ is binary, taking value 0 or 1. In linear regression, testing the null hypothesis $H_0 : \beta_2 = 0$ involves fitting the null model (i.e. $X_1$ only; we take the intercept as given), and the alternative model (i.e. $X_1$ and $X_2$). Then $H_0$ is tested by comparing the sums of squared residuals under both models in an $F$-test. In distributional regression, we test the null hypothesis $H_0 : Q_\oplus(x_1, x_2) = Q_\oplus(x_1)$, that is, $X_2$ does not influence the conditional mean distribution. We fit the null model and the alternative model, and compare the relative squared metric distances between observations and their respective conditional mean curves [2].

### S2 P-value Correction

#### S2.1 Misspecified Null Distribution

In the general regression setting, suppose we have test statistic $T \equiv T(X)$, where large values indicate evidence against the null hypothesis $H_0 : T \sim F_0$. For simplicity, let $F_0$ be absolutely continuous with respect to the Lebesgue measure ("a.c."). The $\alpha$-level hypothesis test for $H_0$ is thus to reject iff

$$T \geq Q_{F_0}(1-\alpha) \qquad \Longleftrightarrow \qquad P := 1 - F_0(T) \leq \alpha.$$

If indeed $T \sim F_0$ under $H_0$, then the CDF of $P$ under $H_0$, which we denote $F_{P,0}$, is

$$\begin{aligned} F_{P,0}(u) &= \mathbf{P}[P \leq u | T \sim F_0] \\ &= \mathbf{P}[1 - F_0(T) \leq u | T \sim F_0] \\ &= \mathbf{P}[T \geq F_0^{-1}(1-u) | T \sim F_0] \\ &= 1 - (F_0 \circ F_0^{-1})(1-u) \\ &= u, \end{aligned}$$

which is the CDF of the standard uniform distribution. However, if $F_0$ is a misspecification, and in reality $H_0 : T \sim G_0 \neq F_0$ (again for simplification, $G_0$ is a.c.), then

$$F_{P,0}(u) = 1 - (G_0 \circ F_0^{-1})(1-u). \tag{S5}$$

Since $G_0 \circ F_0^{-1}$ is not the identity map, we do not obtain the cancellation as in the previous case, and $F_{P,0} \neq \text{unif}(0,1)$. The test $P \leq \alpha$ does not have size $\alpha$, and we must correct $P$.

#### S2.2 Null-Corrected P-value

The p-value $P := 1 - F_0(T)$ of a test statistic $T \equiv T(X)$ is itself a test statistic, being a function only of data $X$ and parameters of a *putative* distribution for $T$, and having a rejection region $P \leq \alpha$. When the null distribution $T \sim F_0$ is correctly specified, $\mathbf{P}[P \leq \alpha] = \alpha$, and $P \leq \alpha$ is immediately an $\alpha$-size statistical test for $H_0$. However, if $T \sim F_0$ is an incorrect specification under $H_0$, then $F_{P,0} \neq \text{unif}(0,1)$, and $P \leq \alpha$ is not an $\alpha$-size test.

If $P \sim F_{P,0}$ under $H_0$ (and $F_{P,0}$ is a.c.), note

$$\begin{aligned} \mathbf{P}[F_{P,0}(P) \leq u | P \sim F_{P,0}] &= \mathbf{P}[P \leq F_{P,0}^{-1}(u) | P \sim F_{P,0}] \\ &= (F_{P,0} \circ F_{P,0}^{-1})(u) \\ &= u. \end{aligned}$$

Hence

$$P_C := F_{P,0}(P) \sim \text{unif}(0,1), \tag{S6}$$

and (since cumulative distribution functions are monotone non-decreasing) $P_C \leq \alpha$ is an $\alpha$-size test for $H_0$.

#### S2.3 Null-Corrected Power

The power of a statistical test is defined as the probability of rejecting $H_0$ with a specific test statistic, assuming a competing alternative hypothesis $H_a$ is true. Using $P_C \leq \alpha$ as a statistical test of appropriate size, we posit alternative $H_a : P \sim F_{P,a} \neq F_{P,0}$, where $F_{P,a}$ is stochastically dominated by $F_{P,0}$ (i.e. $F_{P,a} \geq F_{P,0}$, with strict inequality on some set of non-zero measure). The corrected power is

$$\begin{aligned} \text{power}_{H_a}(\alpha) &:= \mathbf{P}[P_C \leq \alpha | P \sim F_{P,a}] \\ &= \mathbf{P}[F_{P,0}(P) \leq \alpha | P \sim F_{P,a}] \\ &= \mathbf{P}[P \leq F_{P,0}^{-1}(\alpha) | P \sim F_{P,a}] \\ &= (F_{P,a} \circ F_{P,0}^{-1})(\alpha) \geq \alpha, \end{aligned} \tag{S7}$$

where the last inequality follows from the stochastic dominance property.

#### S2.4 Applying Corrections in Practice

If $F_{P,0}$ is analytically known, calculating $P_C = F_{P,0}(P)$ is trivial. In the case the null p-value distribution is not known, but we can obtain samples from it, we can correct the test p-values (and correct the estimated power) utilizing the empirical CDF $\widehat{F}_{P,0}$.

##### S2.4.1 Empirical P-value Correction

Suppose $P \sim F_P$, and let $P_{k,0} \overset{\text{iid}}{\sim} F_{P,0}$ (not necessarily the same as $F_P$) for $k = 1, \ldots, K$. The empirically corrected p-value is given by

$$\begin{aligned} P_C &:= F_{P,0}(P) \\ &\approx \widehat{F}_{P,0}(P) \\ \widehat{P}_C &= \frac{1}{K} \sum_{k=1}^{K} \mathbb{1}\{P_{k,0} \leq P\}, \end{aligned}$$

where $\widehat{P}_C \overset{\text{a.s.}}{\to} F_{P,0}(P) = P_C$ by the strong law of large numbers. Note, however, this procedure is not feasible for very small values of $P$, as precisely estimating $P_C$ (i.e. distinguishing $\widehat{P}_C$ from zero via small standard error) requires a correspondingly very large number of samples.

##### S2.4.2 Empirical Power Correction

Suppose $H_a : P \sim F_{P,a}$ is an alternative hypothesis to $H_0 : P \sim F_{P,0}$, where $F_{P,0}$ stochastically dominates $F_{P,a}$. Let $P_{k,0} \overset{\text{iid}}{\sim} F_{P,0}$ for $k = 1, \ldots, K$, and let $P_\ell \overset{\text{iid}}{\sim} F_{P,a}$ for $\ell = 1, \ldots, L$. Then estimating the corrected power (S7) for $\alpha$-level test $P_C \leq \alpha$ is accomplished by

$$\begin{aligned} \text{power}_{H_a}(\alpha) &= (F_{P,a} \circ F_{P,0}^{-1})(\alpha) \\ &\approx \frac{1}{L} \sum_{\ell=1}^{L} \mathbb{1}\{P_\ell \leq \widehat{F}_{P,0}^{-1}(\alpha)\} \\ \widehat{\text{power}}_{H_a}(\alpha) &= \frac{1}{L} \sum_{\ell=1}^{L} \mathbb{1}\left\{P_\ell \leq P_{(\gamma),0}\right\}, \qquad \gamma := \lceil \alpha K \rceil, \end{aligned}$$

where $P_{(\gamma),0}$ is the $\gamma^{\text{th}}$ order statistic of the $P_{k,0}$'s.

#### S3 Figures

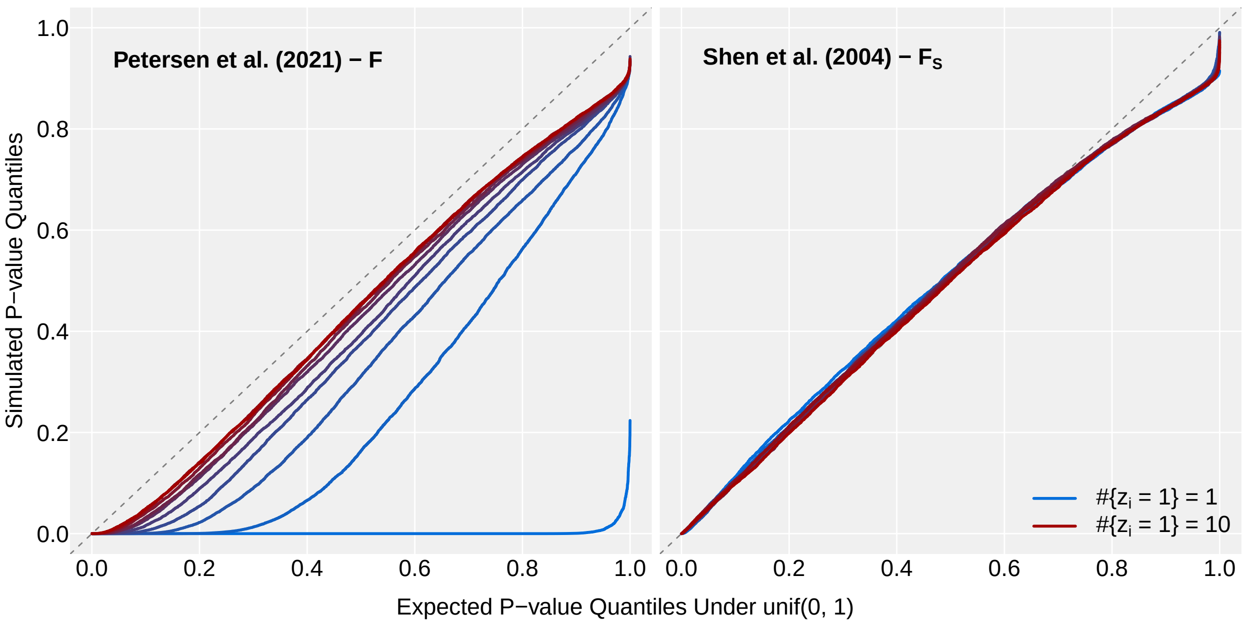


**Figure S1:** QQ-plots of simulated partial $F$-test p-values against expected quantiles under $\text{unif}(0, 1)$, for differing number of "rare allele" observations $|\mathcal{I}| = 1, 2, \ldots, 10$ among 1000 samples. Results across 10,000 iterations each setting.

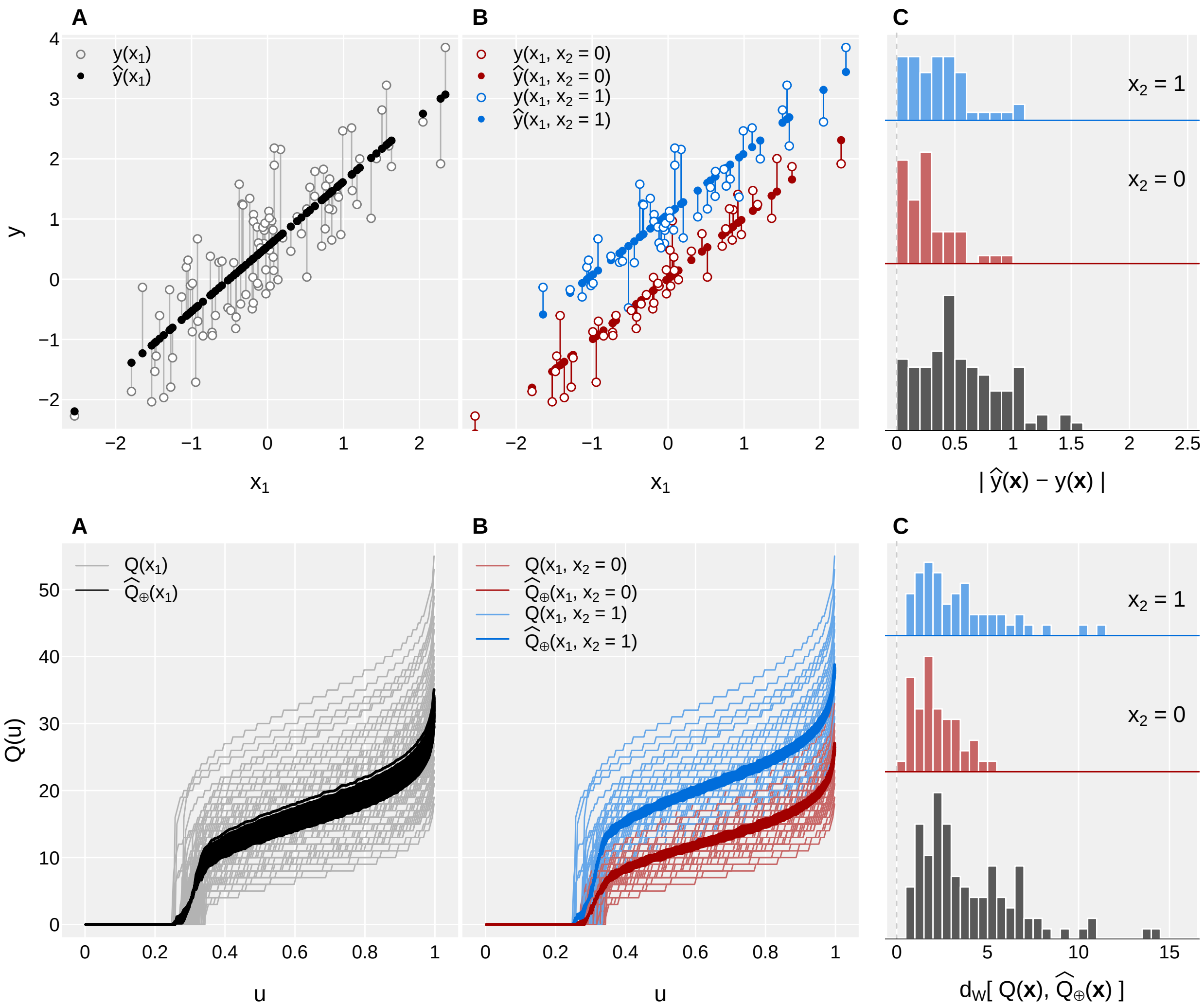


**Figure S2:** Hypothesis testing in linear regression (*top row*) and distributional regression (*bottom row*). Responses are dependent on $X_1$ and $X_2$, where $X_2$ is binary 0 or 1. Testing null hypothesis $H_0 : \beta_2 = 0$ (linear) or $H_0 : Q_\oplus(X_1, X_2) = Q_\oplus(X_1)$ (distributional) involves fitting the null model, i.e. $X_1$ only (panels **A**); fitting the alternative model, i.e. including $X_2$ (panels **B**); and comparing distances between observed and fitted values (panels **C**) with an F test. In each panel **C**, null model distances are shown in the lower histogram, and alternative model distances are shown in the middle (cases where $X_2 = 0$) and upper (cases where $X_2 = 1$).

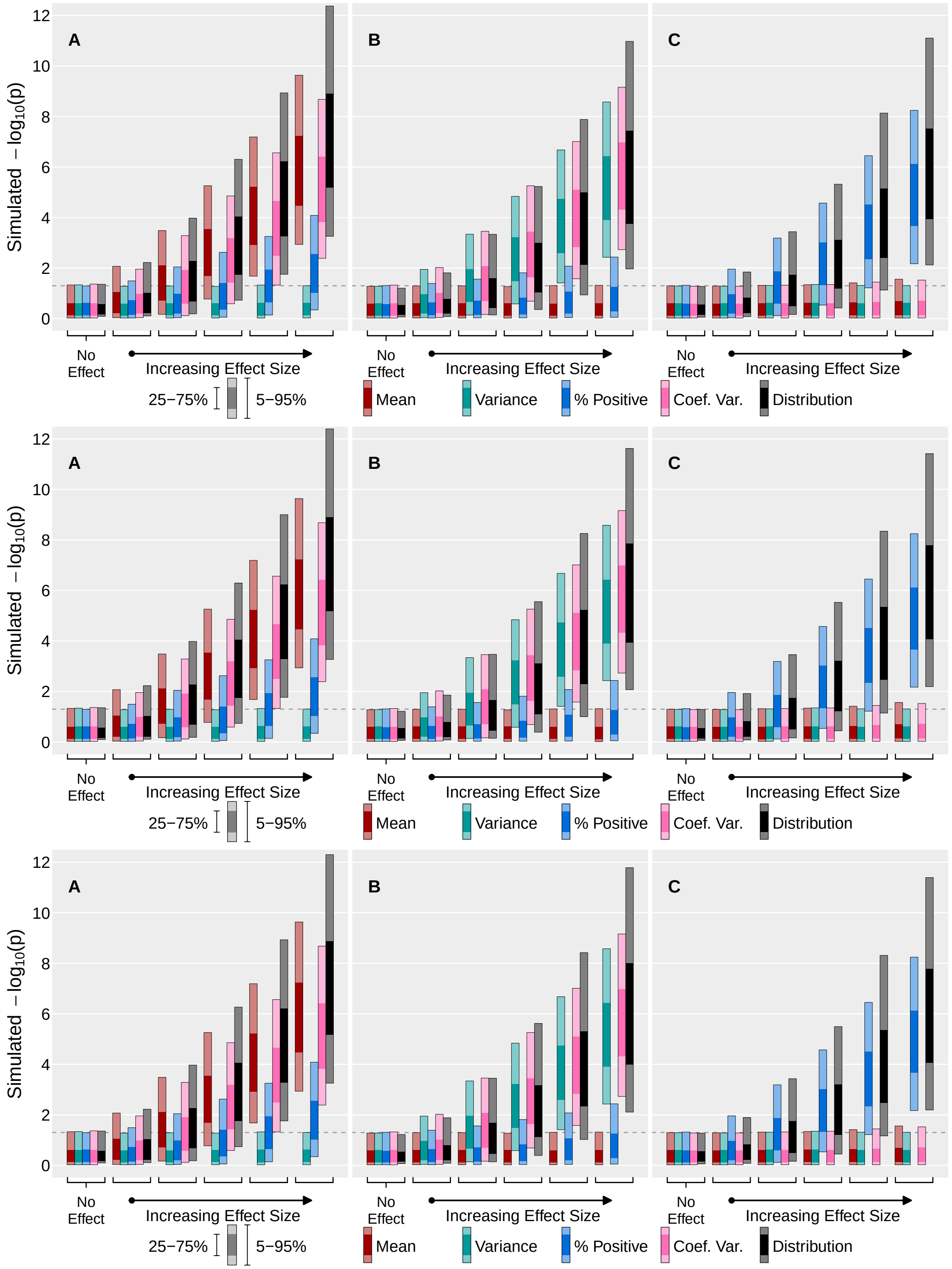


**Figure S3:** Comparison of $\mu$QTL, vQTL, bQTL (% positive/burst), cvQTL, and distQTL test of $X_1$ on simulated ZINB data, with sample size $n = 100$, and $p = 5$ covariates, and $N \sim \text{Poisson}(200)$ observations per sample. **A**: $X_1$ influences mean only, not variance; **B**: $X_1$ influences variance only, not mean; **C**: $X_1$ influences $\text{P}[Y = 0]$ only, not mean or variance. **Top row**: $m = 100$ distQTL quantile discretization; **Middle row**: $m = 150$ distQTL quantile discretization; **Bottom row**: $m = 200$ distQTL quantile discretization. Results over 5000 iterations for each setting.

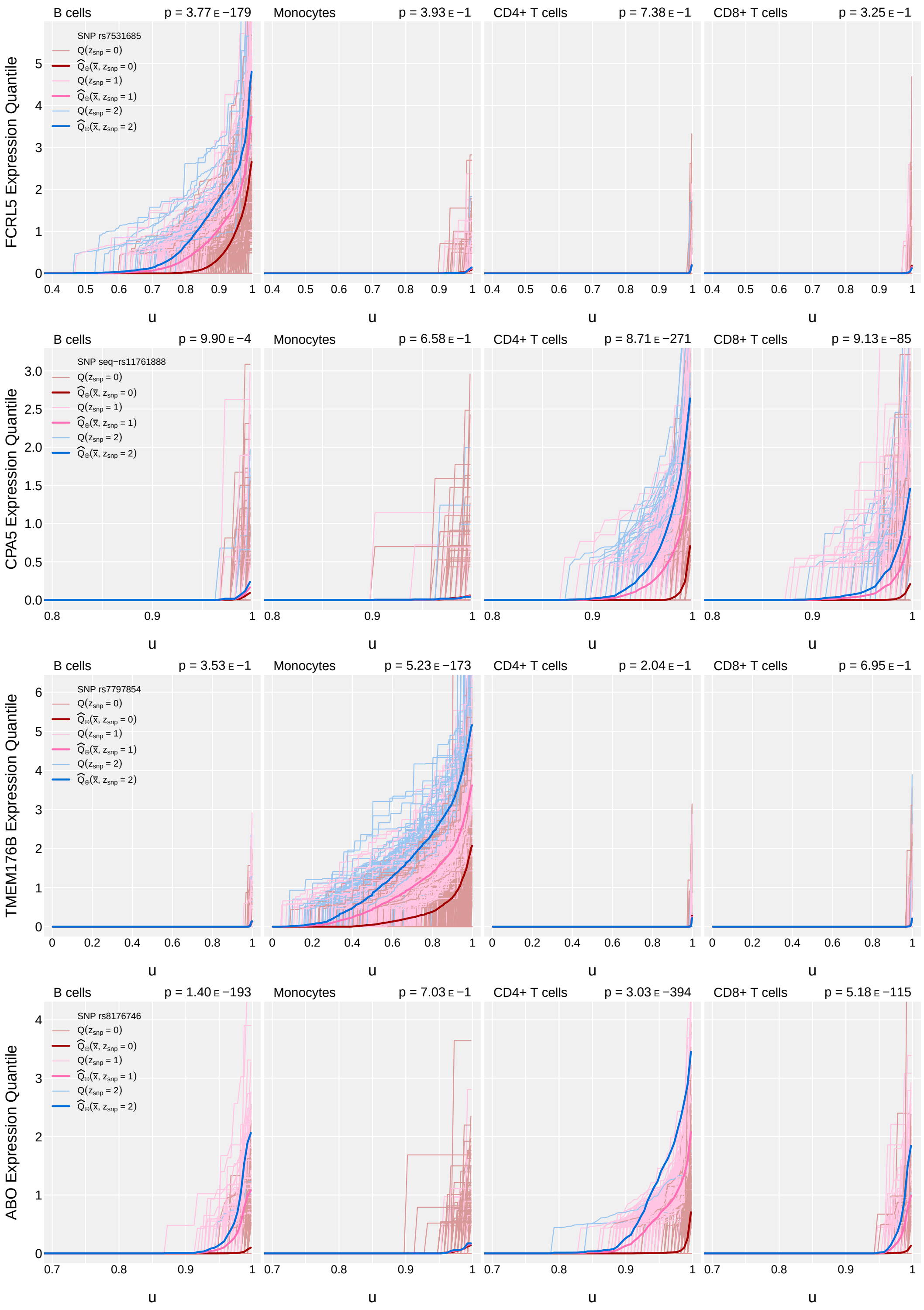


**Figure S4:** Example gene-SNP pairs with differential expression across cell type groups, as identified by distQTL. Raw distQTL p-values are reported since not all genes are “high expressing”.

(A)

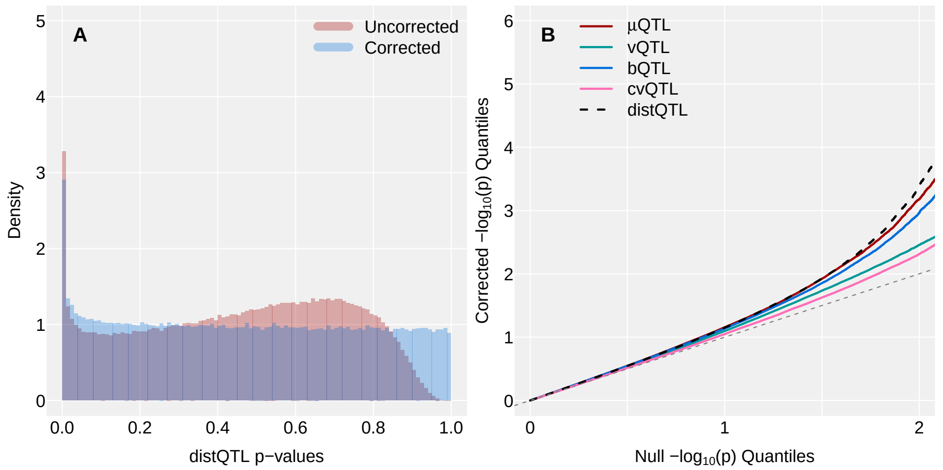


(B)

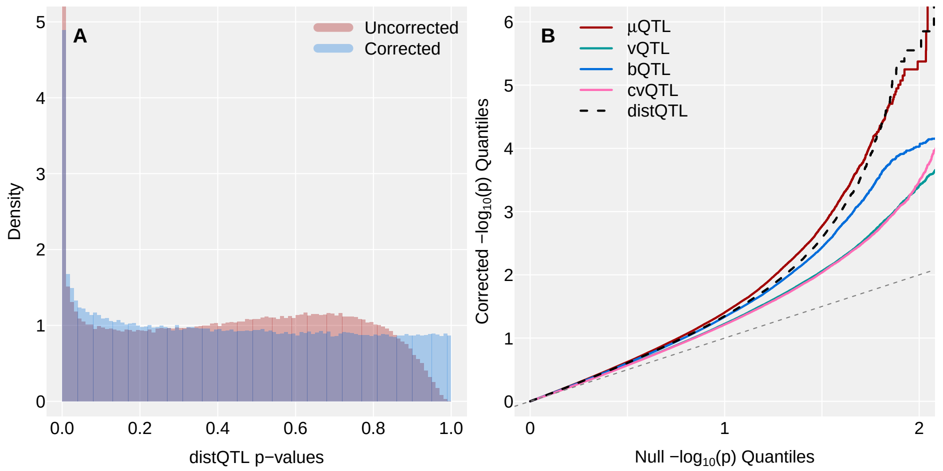


(C)

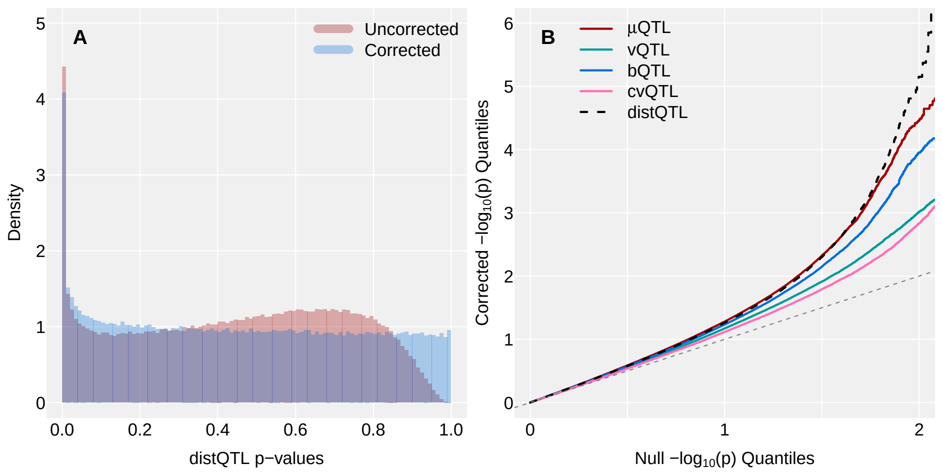


**Figure S5:** P-value correction. Left panels: empirical histograms of raw and null-corrected distQTL p-values from gene-SNP pair models in (A) monocytes, (B) CD4+ T cells, and (C) CD8+ T cells. Right panels: QQ-plots of null-corrected $-\log_{10}(p)$-values for gene-SNP pairs.

(A)

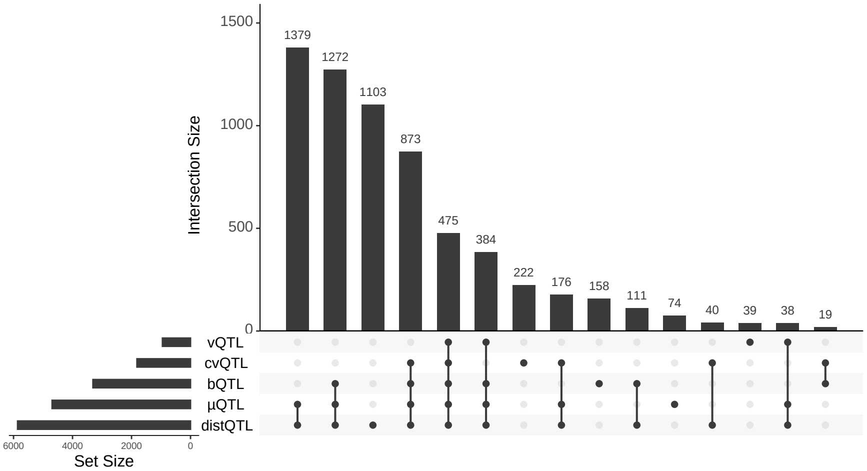


(B)

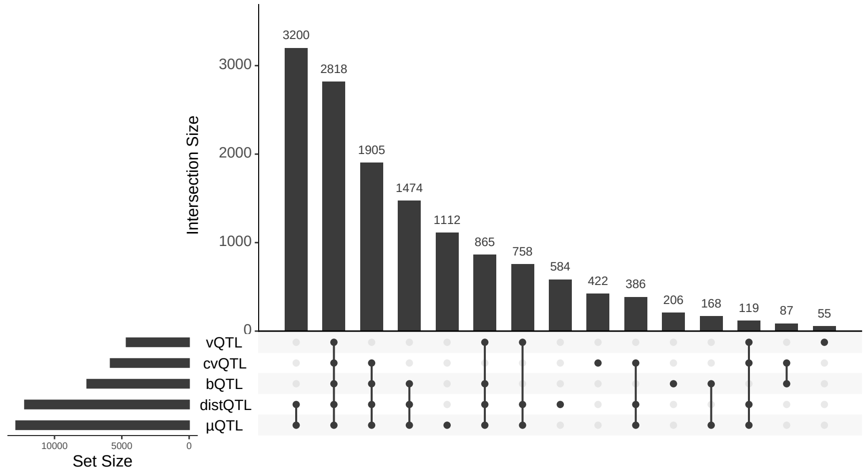


(C)

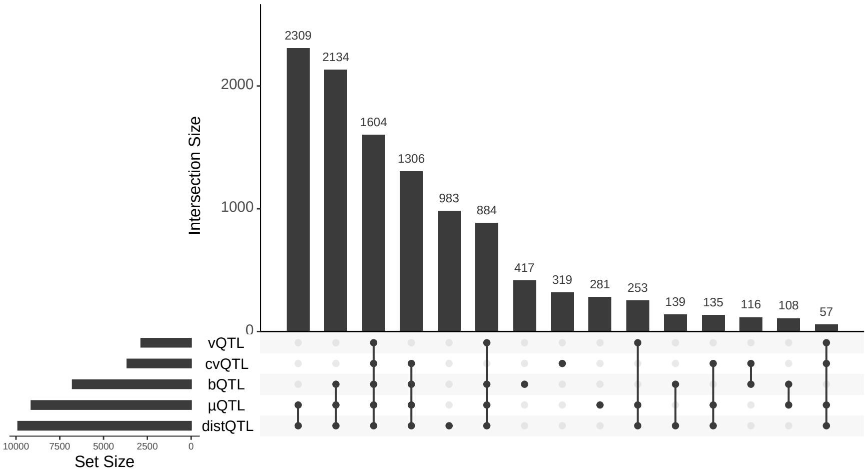


**Figure S6:** Upset plot of eQTL-labeled gene-SNP pairs across (A) monocytes, (B) CD4+ T cells, and (C) CD8+ T cells, using $\alpha = 10^{-4}$ significance threshold. Truncated to 15 intersections.

(A) B cells

| | | When this method is *n.s.* ($p > 0.1$) | | | | |
|---|---|---|---|---|---|---|
| | | $\mu$QTL | vQTL | bQTL | cvQTL | distQTL |
| When this method is significant ($p \leq 10^{-4}$) | $\mu$QTL | · | 148 | 403 | 1991 | 0 |
| | vQTL | 5 | · | 109 | 508 | 0 |
| | bQTL | 37 | 36 | · | 65 | 37 |
| | cvQTL | 45 | 217 | 36 | · | 25 |
| | distQTL | 4 | 163 | 532 | 2409 | · |

(B) Monocytes

| | | When this method is *n.s.* ($p > 0.1$) | | | | |
|---|---|---|---|---|---|---|
| | | $\mu$QTL | vQTL | bQTL | cvQTL | distQTL |
| When this method is significant ($p \leq 10^{-4}$) | $\mu$QTL | · | 40 | 161 | 1246 | 0 |
| | vQTL | 2 | · | 52 | 207 | 0 |
| | bQTL | 56 | 107 | · | 965 | 53 |
| | cvQTL | 75 | 146 | 56 | · | 28 |
| | distQTL | 1 | 62 | 247 | 1850 | · |

(C) CD4+ T cells

| | | When this method is *n.s.* ($p > 0.1$) | | | | |
|---|---|---|---|---|---|---|
| | | $\mu$QTL | vQTL | bQTL | cvQTL | distQTL |
| When this method is significant ($p \leq 10^{-4}$) | $\mu$QTL | · | 339 | 482 | 1495 | 0 |
| | vQTL | 11 | · | 143 | 425 | 1 |
| | bQTL | 39 | 225 | · | 661 | 37 |
| | cvQTL | 33 | 403 | 17 | · | 13 |
| | distQTL | 5 | 240 | 547 | 1503 | · |

(D) CD8+ T cells

| | | When this method is *n.s.* ($p > 0.1$) | | | | |
|---|---|---|---|---|---|---|
| | | $\mu$QTL | vQTL | bQTL | cvQTL | distQTL |
| When this method is significant ($p \leq 10^{-4}$) | $\mu$QTL | · | 180 | 296 | 1999 | 0 |
| | vQTL | 2 | · | 42 | 495 | 4 |
| | bQTL | 26 | 253 | · | 1554 | 32 |
| | cvQTL | 68 | 285 | 51 | · | 43 |
| | distQTL | 3 | 179 | 403 | 2426 | · |

**Table S1:** Cross-tabulation of counts of gene-SNP pairs where one eQTL-mapping method identifies a SNP effect as very significant (corrected $p \leq 10^{-4}$) while another identifies it as very non-significant (*n.s.*, corrected $p > 0.1$). Large counts in a method's row indicate it identifies more eQTLs while other methods do not; large counts in a method's column indicate it does not identify eQTLs that other methods do. Results are shown for (A) B cells, (B) monocytes, (C) CD4+ T cells, and (D) CD8+ T cells separately.

| | | | $\log_{10}(p)$ | | | | |
|---|---|---|---|---|---|---|---|
| Group | Gene ENSID | SNP | $\mu$QTL | vQTL | bQTL | cvQTL | distQTL |
| B | ENSG00000115350 | rs1861457 | -0.042 | -2.960 | -2.844 | -3.948 | -4.705 |
| B | ENSG00000115350 | rs12713829 | -0.153 | -2.600 | -2.853 | -4.088 | -4.333 |
| B | ENSG00000115350 | rs6741029 | -0.270 | -2.461 | -2.968 | -4.066 | -4.218 |
| B | ENSG00000164144 | GSA-rs114228794 | -0.860 | -1.675 | -0.005 | -2.495 | -4.073 |
| Mono | ENSG00000188536 | GSA-rs190072257 | -0.098 | -3.558 | -0.550 | -0.582 | -4.158 |
| CD4T | ENSG00000136521 | GSA-rs35399127 | -0.386 | -3.826 | -1.197 | -1.638 | -4.170 |
| CD4T | ENSG00000100941 | chr14-39594241 | -0.263 | -3.959 | -3.469 | -4.319 | -5.550 |
| CD4T | ENSG00000128463 | GSA-rs75544400 | -0.112 | -2.793 | -2.294 | -1.725 | -4.198 |
| CD4T | ENSG00000185905 | rs7193756 | -0.085 | -3.232 | -4.198 | -5.851 | -5.073 |
| CD4T | ENSG00000091592 | GSA-rs117883712 | -0.229 | -1.773 | -1.973 | -3.020 | -5.374 |
| CD8T | ENSG00000198502 | rs144757939 | -0.791 | -1.348 | -0.312 | -0.819 | -9.326 |
| CD8T | ENSG00000182400 | chr14-39594241 | -0.986 | -3.122 | -0.341 | -1.492 | -4.619 |
| CD8T | ENSG00000185905 | rs7193756 | -0.678 | -3.585 | -3.021 | -4.134 | -4.646 |
| All | ENSG00000138363 | rs4673980 | -0.618 | -3.434 | -1.042 | -0.668 | -4.024 |
| All | ENSG00000123200 | GSA-rs182108787 | -0.204 | -1.919 | -1.907 | -3.506 | -4.252 |
| All | ENSG00000182400 | chr14-39594241 | -0.364 | -3.694 | -2.616 | -2.505 | -5.875 |
| All | ENSG00000150527 | chr14-39594241 | -0.263 | -2.697 | -4.645 | -7.105 | -5.176 |
| All | ENSG00000117906 | GSA-rs117151724 | -0.008 | -2.513 | -3.010 | -4.514 | -4.796 |
| All | ENSG00000185905 | rs7193756 | -0.043 | -4.263 | -8.614 | -11.026 | -8.181 |
| All | ENSG00000091640 | rs6502844 | -0.840 | -1.155 | -4.671 | -4.090 | -4.252 |
| All | ENSG00000263753 | GSA-rs8082898 | -0.594 | -3.775 | -0.775 | -1.068 | -4.062 |
| All | ENSG00000101365 | rs4815541 | -0.127 | -2.934 | -2.367 | -2.118 | -4.024 |
| All | ENSG00000211679 | rs2519501 | -0.863 | -4.796 | -0.527 | -3.807 | -9.904 |
| All | ENSG00000211679 | rs737885 | -0.755 | -3.357 | -1.027 | -3.285 | -4.796 |
| All | ENSG00000100412 | GSA-rs11704973 | -0.116 | -3.170 | -2.077 | -3.066 | -4.972 |

**Table S2:** Cell-type-specific gene-SNP pairs where distQTL identifies a SNP effect as significant (corrected $p \leq 10^{-4}$) while $\mu$QTL identifies it as very non-significant (*n.s.*; corrected $p > 0.10$). Of note, there are no corresponding gene-SNP pairs where $\mu$QTL is significant at $p \leq 10^{-4}$ and distQTL is *n.s.* at $p > 0.1$.

(A)

| **Cell group:** | B | Mono | CD4+ T | CD8+ T |
|---|---|---|---|---|
| OneK1K % | 20.0 | 8.3 | 43.4 | 28.2 |

(B)

| **Cell group:** | B | Mono | CD4+ T | CD8+ T |
|---|---|---|---|---|
| $\mu$QTL | 12.9 | 8.3 | 23.3 | 15.1 |
| vQTL | 4.3 | 1.8 | 9.5 | 5.3 |
| bQTL | 0.5 | 6.4 | 16.6 | 12.9 |
| cvQTL | 5.5 | 3.5 | 13.0 | 7.1 |
| distQTL | 13.7 | 9.9 | 22.2 | 16.1 |

(C)

| **Cell group:** | B | Mono | CD4+ T | CD8+ T |
|---|---|---|---|---|
| $\mu$QTL | 12.4 | 8.2 | 19.0 | 14.0 |
| vQTL | 4.9 | 2.0 | 7.8 | 4.9 |
| bQTL | 0.6 | 6.2 | 12.1 | 11.1 |
| cvQTL | 5.2 | 3.0 | 8.9 | 5.2 |
| distQTL | 13.4 | 10.2 | 18.1 | 14.7 |

**Table S3:** eQTL-mapping overlap between GTEx and cell-type-specific analysis in OneK1K cohort is higher for cell types with higher relative proportion in blood samples. (A) Percentage of cells in post-QC OneK1K cohort from each cell-type group; percentages may not add to 100% due to rounding. (B) Percentage GTEx eGenes recovered from OneK1K cell-type-specific analysis. (C) Percentage GTEx eQTL recovered from OneK1K cell type specific analysis. eQTL identification is made for a gene-SNP pair in both GTEx and OneK1K if corrected (as applicable) p-value $p \leq 10^{-4}$.

### References

[1] Petersen, A., Liu, X., and Divani, A. A. Wasserstein F-tests and confidence bands for the Fréchet regression of density response curves. Ann. Stat. 2021; 49(1): 590–611.

[2] Shen, Q. and Faraway, J. An F Test for Linear Models with Functional Responses. Stat. Sin. 2004; 14(4): 1239–1257.
